# Leishmania PNUTS discriminates between PP1 catalytic subunits through a RVxF-ΦΦ-F motif and polymorphisms in the PP1 C-tail and catalytic domain

**DOI:** 10.1101/2023.09.20.558696

**Authors:** Yang Zhang, Robert Sabatini

## Abstract

PP1 phosphatases lack substrate specificity and associate with specific regulatory subunits to achieve selectivity. Among the eight PP1 isotypes in Leishmania, PP1-8e associates with the regulatory protein PNUTS along with the structural factors JBP3 and Wdr82 in the PJW/PP1 complex that modulates RNA polymerase II (Pol II) phosphorylation and transcription termination. Little is known regarding interactions involved in PJW/PP1 complex formation, including how PP1-8e is the selective isotype associated with PNUTS. Here, we show that PNUTS uses an established RVxF-ΦΦ-F motif to bind the PP1 catalytic domain with similar interfacial interactions as mammalian PP1- PNUTS and non-canonical motifs. These atypical interactions involve residues within the PP1-8e catalytic domain and N- and C-terminus for isoform specific regulator binding. This work advances our understanding of PP1 isoform selectivity and reveals key roles of PP1 residues in regulator binding. We also explore the role of PNUTS as a scaffold protein for the complex by identifying the C-terminal region involved in binding JBP3 and Wdr82, and impact of PNUTS on the stability of complex components and function in Pol II transcription *in vivo*. Taken together, these studies provide a potential mechanism where multiple motifs within PNUTS are used combinatorially to tune binding affinity to PP1, and the C-termini for independent binding of JBP3 and Wdr82, in the Leishmania PJW/PP1 complex. Overall, our data provide insights in the formation of the PJW/PP1 complex involved in regulating Pol II transcription in divergent protozoans where little is understood.

## Introduction

Phosphorylation is a critical regulatory mechanism for over 70% of eukaryotic cellular proteins, and the majority of the phosphorylations occur on serine, threonine or tyrosine residues (1). More than 420 serine/threonine kinases target specific serine/threonine residues, which account for approximately 98% of all phosphorylation events. On the other hand, fewer than 40 serine/threonine phosphatases are involved in protein dephosphorylation (2, 3). Protein Phosphatase 1 (PP1) is a major serine/threonine phosphatase, estimated to catalyze one third of all dephosphorylation events in eukaryotic cells and involved in many essential cellular activities (including cardiac muscle contraction, glycogen metabolism, cell cycle transition, and transcription termination)(4–6). In contrast to protein serine/threonine kinases, PP1 has little intrinsic substrate specificity. To carry out specific functions in a wide variety of cellular activities, PP1 binds over 200 confirmed PP1-interacting proteins (PIPs), forming highly specific holoenzymes in mammalian cells (3). These PIPs target PP1 to distinct cellular compartments and/or direct its activity toward specific substrates (7). PIPs usually associate with PP1 using a combination of short linear motifs (SLiMs). They bind in a largely extended manner at multiple sites across the top of PP1 (remote from the catalytic site), including the RVxF motif binding site and the ΦΦ motif binding site, both of which are used by a large number of PIPs. However, many studies have shown that PIP binding is usually more complex, with PIPs utilizing additional motifs beyond the RvXF and ΦΦ motif for PP1 holoenzyme formation (8). Characterizing these interactions is key to understanding how PIPs associate with PP1 and regulate specific biological processes such as transcription and gene expression.

One of the earliest characterized PIPs is PNUTS (PP1 nuclear targeting subunit), originally described as a nuclear regulator of PP1 that helps retain PP1 in the nucleus (9–11). PNUTS has been implicated in PP1-regulated processes including cell cycle regulation (12), RNA processing (9, 13), DNA repair (14), transcription (15) and telomere stability (16). Like most PIPs, PNUTS is a largely unstructured protein in the unbound state and included in a group of intrinsically disordered proteins (IDPs) (7). This intrinsic flexibility is important for the formation of extensive interactions with PP1. PNUTS modulation of PP1 is mediated by a central region, employing RVxF-ΦΦ-Phe-Arg motifs (17). The most well-characterized motif is the RVxF motif ([K/R]-X_0–1_-[V/I/L]-X-[F/W], where X can be any amino acid except proline) that is found in 90% of PIPs (18–20). ^398^TVTW^401^ in human PNUTS (hPNUTS) constitutes the canonical RVxF PP1-binding motif, with the second and fourth residues burying deep in two hydrophobic pockets on the PP1 surface, providing an essential stabilizing force (17). As demonstrated for hPNUTS (17, 21), mutation of hydrophobic valine and phenylalanine/tryptophan positions in the RVXF-binding motif typically abolishes the ability of PIP to bind to PP1.

Structure analyses of PIP:PP1 holoenzymes (including PNUTS) have identified several additional motifs that make contact with PP1(17). For example, the ΦΦ motif is a two-hydrophobic residue motif that is usually found 5-8 amino acids C-terminal to the RVXF motif on PIPs (17). hPNUTS-PP1 is found to be associated with two additional structural proteins, Wdr82 and the DNA binding protein Tox4, in a complex called PTW/PP1 (21). PNUTS is the scaffolding protein in the complex and mediates independent associations of PP1, Wdr82 and Tox4. Tox4 interacts with an N-terminal TFIIS domain in hPNUTS, while Wdr82 binds to a C-terminal region in hPNUTS (aa 418-619). The PTW/PP1 complex is a negative regulator of RNA Pol II elongation rate and plays a key role in transcription termination.

Depletion of individual components in human cells, or ortholog components in yeast, leads to RNA Pol II transcription termination defects (15, 22–25). In the torpedo model of transcription termination, as Pol II reaches the poly(A) signal, pre-mRNA is cleaved, providing an entry site for the 5’-3’ exoribonuclease Xrn2 to catch up with the Pol II and dislodge it from the DNA template, allowing for transcription termination (26–28). Dephosphorylation of the Pol II C-terminal domain (CTD) and Spt5, reducing the speed of the polymerase within the so-called termination zone, facilitates this process (29–31).

The Trypanosomatidae are early divergent protozoan parasites. Several members of the Trypanosomatidae including *Trypanosoma brucei* and *Leishmania major* are pathogenic to humans, causing Human African Trypanosomiasis (African sleeping sickness) and leishmaniasis. In these parasites, hundreds of genes of unrelated functions are arranged into polycistronic transcription units (PTUs) throughout the genome (32, 33). Genes in each PTU are co-transcribed from an initiation site at the 5’ end to the termination site at the 3’ end. Pre-mRNAs are processed through trans-splicing with the addition of a 39-nucleotide spliced leader sequence to the 5’ end of mRNAs, which is coupled to the 3’ polyadenylation of the upstream transcript (34–41). Very little is understood regarding the RNA Pol II transcription cycle (initiation, elongation and termination) in these important eukaryotic pathogens.

Epigenetic markers, such as histone variants (H3V and H4V) and the DNA modification base J, are enriched at Pol II transcription termination sites in Leishmania and *T. brucei* (32, 42–44). Base J is a glucosylated thymidine (45) and has only been identified in the nuclear DNA of kinetoplastids, *Diplonema,* and *Euglena* (46, 47). The loss of base J (and H3V) in Leishmania and *T. brucei* led to readthrough transcription at termination sites, suggesting a critical role of base J in Pol II transcription termination (48–52). Exploring base J function further led to the identification of the PJW/PP1 complex in *Leishmania tarentolae* composed of PP1-PNUTS-Wdr82 and a base J-binding protein, JBP3 (53, 54).

LtPNUTS is a predictively disordered 29 kDa protein with 23% sequence identity to hPNUTS, and contains a putative RVxF PP1 binding motif (^97^RVCW^99^) (53). Alanine substitution of the hydrophobic residues in the RVxF motif (^97^RACA^99^) has been shown to disrupt LtPNUTS-PP1 association (55). Additionally, short synthetic RVXF-containing peptides are sufficient to disrupt the LtPNUTS-PP1 association. Ablation of PNUTS, JBP3 and Wdr82 by RNAi in *T. brucei* (53), and deletion of PP1-8e and JBP3 in Leishmania (54, 55), has been shown to cause Pol II termination defects, similar to the defects following the loss of base J/H3V. These *in vivo* data, along with the recent demonstration that Pol II is a direct substrate for PP1-8e as a component of the Leishmania PJW/PP1 complex *in vitro* (55), supports a conserved PNUTS-PP1 regulatory mechanism from trypanosomatids to yeast and mammalian cells. We therefore proposed that similar to the PTW/PP1 complex, LtPNUTS is scaffolding protein that mediates independent binding of PP1, JBP3 and Wdr82, with JBP3 tethering the complex to the base J-enriched transcription termination sites for PP1- mediated dephosphorylation of Pol II.

Eight PP1 isoforms, grouped into five different clades (A-E), are identified in the Leishmania genome (Fig. S1).

Among these, only PP1-8e is found associated with the PJW/PP1 complex *in vivo* and shown to be involved in Pol II transcription termination (55). Although the *T. brucei* genome also harbors eight PP1 isoforms, no obvious PP1 isoform belongs to clade E as a homologue of LtPP1-8e (55). Furthermore, purification of TbPNUTS pulls down JBP3 and Wdr82 but not PP1(53). Presumably, transient/weak association between a TbPP1 isotype and the PNUTS-Wdr-JBP3 complex via the conserved RVxF PP1-binding motif, allows a conserved transcription termination mechanism in *T. brucei* cells (53, 55). Unique sequences within PP1-8e may explain isotype selectivity of PNUTS binding in Leishmania (55). However, interactions involved in the selectivity of LtPNUTS for the PP1-8e isoform have not been explored. In fact, while PP1 isoform selectivity is thought to be an important feature of regulatory PIPs, limited mechanistic information exists on how this is achieved in any system. The mammalian PP1 isoforms (PP1a, PP1b, PP1g) share a sequence identity ranging from 85% to 93%, and sequence variability mainly comes from the divergent N- and, most notably, C-termini, with only a few amino acid residues being different within the catalytic domains (2). Among the regulatory PIPs which display isoform preferences, such as MYPT1(56, 57), Spinophilin (58), RepoMan (59), Ki67(59), ASPP2 (60) and RRP1B (61), specificity is achieved via recognition of the PP1 C-terminus or a β/γ specificity pocket within the PP1 catalytic domain. The extreme C-terminus of PP1 (PP1α^309-330^) contains a SH3-binding motif (PPII –

PxxPxR), that is conserved among all the mammalian PP1 isoforms, and a variable C-tail. The apoptosis stimulation proteins of p53 family (iASPP/ASPP1/ASPP2) utilize a SH3 domain to selectively bind the PP1 C-terminus via contacts in the PPII motif and residues in the variable C-tail region to achieve isoform selectivity (60, 62, 63). Ankyrin repeats of the myosin phosphatase targeting subunit MYPT1 associate with amino acids in the PP1 C-tail and drive selectivity towards PP1β (2). In the case of RRP1B, RepoMan and Ki-67, the SLiM (KiR or SLIV) immediately downstream of the RVxF motif determines the preference toward PP1γ through a single amino acid change in the catalytic domain of PP1(59, 61, 64). Therefore, isoform specificity is mediated in these PIPs by a single amino acid difference in PP1 at position 20, which is an Arg residue in PP1γ/β and a Gln residue in PP1α.

In this study, we employ structural modeling and mutagenesis analysis to help define how LtPNUTS specifically recruits PP1-8e to the PJW/PP1 complex. First, we confirm that LtPNUTS demonstrates substrate specificity for PP1-8e among the identified LtPP1 isoforms *in vivo* by Co-IP analysis. We show that LtPNUTS binds to PP1 via a combination of well-characterized PP1-interacting motifs including the extended RVxF (RVXF-L_R_-LL) and Phe motif. We also identified unique termini and motifs within LtPP1-8e catalytic domain, including sites not previously shown to bind any PP1 regulator, that are important for PP1-PNUTS interaction. Lastly, we explore the scaffold function of PNUTS by mapping the Wdr82 and JBP3 binding domain at the C-terminus of PNUTS and demonstrate PNUTS protein level is critical for the integrity of the PJW/PP1 complex and function in Pol II termination. Together, these data support a model for extensive interactions between LtPNUTS and PP1-8e and provide key insights into the isoform selectivity of LtPNUTS and its scaffold function in overall stability of the PJW/PP1 complex.

## Results

### LtPNUTS displays PP1 isoform selectivity

Our previous affinity purification-mass spectrometry data (53) indicated that PNUTS is part of a tightly interlinked protein network comprising the PP1 catalytic subunit PP1-8e, JBP3 and Wdr82 in *L. tarentolae* cells. While there are 8 PP1 isotypes in the Leishmania genome (Fig. S1) (54, 55), only PP1-8e was associated with the Leishmania PJW/PP1 complex. In order to understand the specific association of PP1-8e in this complex, we first sought to verify the binary interaction between PNUTS and the *L. tarentoae* PP1 catalytic subunits by co-immunoprecipitation (co-IP) *in vivo*. To do this, we HA-tagged the endogenous loci of LtPNUTS using cas9 and overexpressed various Pd-tagged LtPP1 isotypes from a plasmid (Fig. 1A). LtPP1-3 (LtaPh_3411201) is much smaller (167 aa) than other PP1 isotypes and is predicted to contain a partial catalytic core. We were unable to overexpress Pd-tagged LtPP1-3, suggesting that it could be a truncated pseudogene. Therefore, we analyzed 5 of the 7 complete PP1 isotypes in *L. tarentolae* (representing all 5 clades). Our results show that only PP1-8e can IP a significant fraction of PNUTS, while the other PP1 isotypes (PP1-1, PP1-2, PP1-4 and PP1-7) show no detectable interaction with PNUTS by Co-IP (Fig. 1B). These data confirm the MS analysis of the purified PNUTS-PP1 complex (53, 55) and directly demonstrate for the first time that PNUTS preferentially targets PP1-8e over other isoforms in intact leishmania cells.

**Figure 1.**
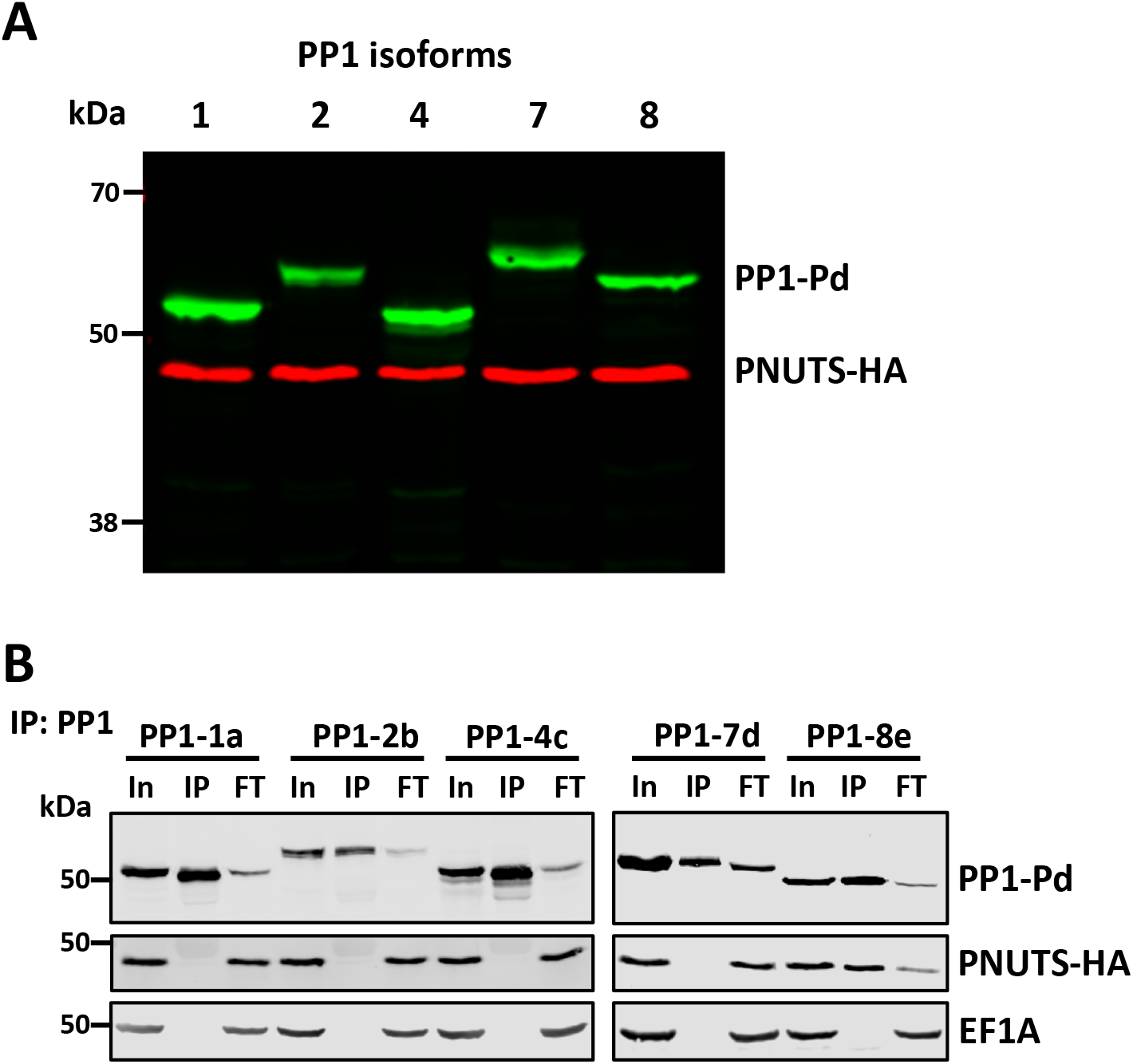
PNUTS binding is specific for the PP1-8e isoform. A, expression of PNUTS and PP1 isoforms in *L. tarentolae*. Cell extracts from *L. tarentolae* cells that endogenously express HA-tagged PNUTS, and exogenously express Pd-tagged PP1 isoforms from the pSNSAP1 vector were analyzed by western blotting with anti-HA and anti-protein A. B, PNUTS/PP1 Co-IP analysis. Lysates from the indicated cell lines were purified by anti-protein A affinity resin and analyzed by western blotting with anti-protein A and anti-HA. Equal cell equivalents of input (In), precipitated immunocomplexes (IP), and flow-through or unbound fraction (FT) were loaded on the gel. EF1A serves as a loading and negative control for the Co-IP.

### Lt PNUTS associates with PP1-8e through an established RVxF-**L_R_-ΦΦ** -F motif

To determine the molecular basis of isoform specificity of PNUTS for PP1-8e in *L. tarentolae*, we used AlphaFold to help define the PNUTS-PP1 interaction interface. We first explored the predicted structure for the LtPP1 isotypes. The PP1 catalytic core is highly conserved across eukaryotes from human to yeast cells, consisting of 10 sets of α-helices (labelled A’ to I) and 15 sets of β-sheets (numbered 1’ to 14) (Fig. S2) (65, 66). The catalytic core regions of the LtPP1 isotypes are predicted to be of high confidence by AlphaFold and their structural overlay to the determined human PP1 protein structure (PDB: 3E7a, Fig. S3A) shows a high structural similarity. An example is the predicted LtPP1-1a structure (Fig. S3B), which shows a high structural identity to hPP1 (Fig. S3C) with a root mean square deviation (RMSD) of 0.580 Å. LtPP1-8e was also predicted with high confidence (Fig. S3D) for the catalytic core region. The predicted LtPP1-8e structure aligns well to the hPP1 structure (Fig. S3E) except three regions within the catalytic core and the N- and C-terminus that we have identified as unique to PP1-8e (55) (Figs. S2 and S3E). Deletion of these unique regions in LtPP1-8e increases the structural similarity between LtPP1-8e and hPP1 (RMSD: 0.666 Å, Fig. S3G). Thus, as previously predicted based on sequence conservation (55), the structural identity of PP1 catalytic subunits between mammals and Leishmania suggest strong functional conservation during evolution. However, unique sequences in PP1-8e may be important for PP1-8e specific functions in Leishmania.

We next submitted LtPNUTS and LtPP1-8e sequences together to AlphaFold2 to generate the predicted LtPP1:PNUTS structure (Fig. S4). As expected, the majority of LtPNUTS is unstructured. While only a limited region of PNUTS is confidently predicted to become buried upon complex formation (Fig. S4), this region binds in a largely extended manner at multiple sites across the top of PP1 in a way highly similar to several well-characterized PP1- interacting proteins (Fig. 2B and C), including human PNUTS (Fig. S5)(17), spinophilin (67), and Gm (68). They share multiple well-charactered PP1-binding motifs, including the RVxF-L_R_-LL binding motif (Fig. 2A and C). Furthermore, similar to hPNUTS, LtPNUTS is predicted to bind PP1 remotely away from the PP1 catalytic site, making it fully accessible to substrate. Consistently, we have recently demonstrated PP1 is catalytically active in the PNUTS:PP1-8e holoenzyme, capable of dephosphorylating model substrates, such as p-nitrophenyl phosphate, as well the LtPol II CTD (55). The first of the key interaction sites in the LtPNUTS:PP1-8e complex is bound by the RVxF-L_R_ motifs (Fig. 2A-D). Nine residues of PNUTS (^93^R to D^101^) adopt an extended conformation and bind to a hydrophobic channel on the PP1 surface formed at the interface of the two β-sheets of the β-sandwich opposite to the catalytic site channel.

**Figure 2.**
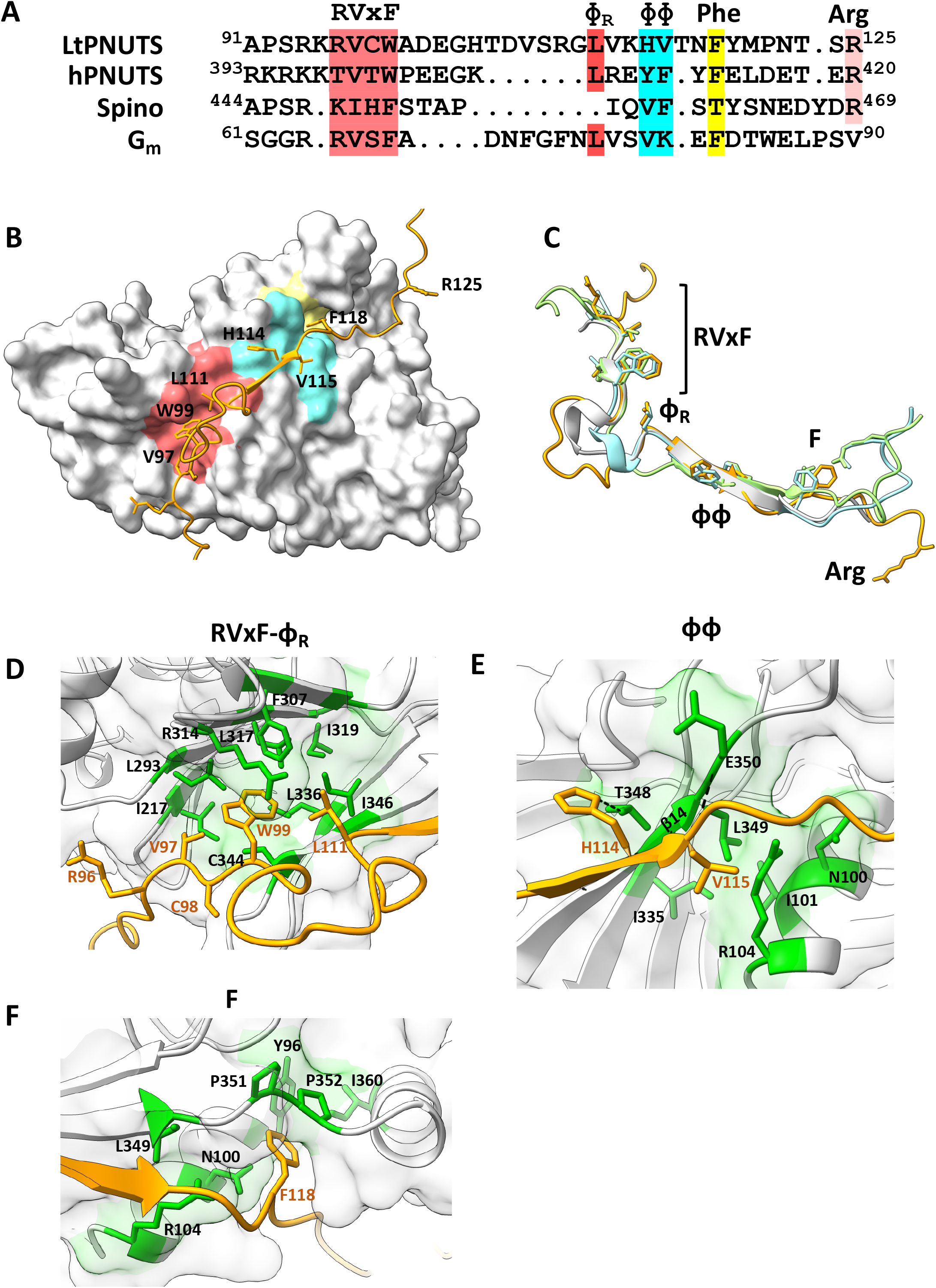
Predicted LtPNUTS-PP1-8e interacting motifs. A, structure-based sequence alignment of the PP1- interacting motifs of LtPNUTS, hPNUTS, spinophilin and Gm, with PP1 interacting residues indicated. B, predicted structure of the LtPNUTS:PP1-8e complex. LtPNUTS is shown as orange ribbon with key interacting residues shown as sticks and LtPP1-8e is shown as a grey surface. LtPNUTS residues ^96^RVCW^99^ and L111 bind to the RVxF binding pocket (red), LtPNUTS residues ^114^HV^115^ bind to the PP1 ΦΦ binding pocket (cyan), and LtPNUTS residues F118 binds to the Phe binding pocket (yellow). The colored regions of PP1-8e correspond to the zoomed in pockets shown in D, E and F. C, overlay of the RVxF and ΦΦ-F structures of four PP1 regulators, LtPNUTS (orange), hPNUTS (green, 4mp0), Spinophilin (blue, 3egg) and Gm (grey, 6dno), with residues binding RVxF-Φ_R_, ΦΦ, and Phe pockets shown as sticks. The Arginine residue (R125) of LtPNUTS that deviates from the hPNUTS and Spinophilin structure is indicated. D-F, major binding interactions between LtPNUTS (orange sticks) and PP1 (surface). The well-established SLiM binding pockets (D, RVxF; E, ΦΦ; F, Phe) are shown. Key interacting residues in LtPP1-8 (black) and LtPNUTS (orange) are labelled. Predicted salt bridge interactions between PP1 residues E350 and T348 with the PNUTS ΦΦ motif indicated by dashed line in E.

PNUTS residues ^96^RVCW^99^ form the RVxF motif, which binds the PP1 RVxF binding pocket, V and W are the anchoring hydrophobic residues that bind deeply in this pocket (Fig. 2D). The predicted LtPNUTS RVxF interaction is highly similar to those observed in other PP1 holoenzyme complexes, including mammalian PNUTS-PP1 (Fig. S5B). Structural and functional studies of the mammalian PP1-PNUTS complex, and modeling of the LtPP1-PNUTS complex here suggest a dominant role for V97 and W99 in stabilizing the interaction between LtPNUTS and PP1-8e. We have recently shown the V97A-W99A double mutant is unable to bind PP1-8e (55). To test this hypothesis in more detail, we made single alanine mutations at each of these positions in the LtPNUTS expression plasmid (pSNSAP1) and tested the PNUTS mutants for interaction by Co-IP with endogenously HA-tagged PP1-8e. Alanine mutation of W99 completely abolished PP1-PNUTS association, and V97A decreased PP1-PNUTS association by 5-fold (Fig. 3A and 3B), indicating the importance of the hydrophobic association mediated by the RVxF motif. Inspection of the structure of hPNUTS in complex with PP1 highlighted interfacial PP1 amino acids I169, L243, F257, R261, V264, I266, M283, C291, F293 that are conserved in LtPP1-8e as I217, L293, F307, R314, L317, L336, V343, C344, and I346 that form the hydrophobic pocket and stabilize V97 and W99 in the PNUTS RVxF motif (Fig. 2D and Fig. S5B). To test this, single alanine mutation of Iso217 was introduced into the LtPP1-8e expression construct and the PP1 mutants tested for interaction by Co-IP with endogenously HA-tagged PNUTS. Mutation of I217_PP1_ to alanine significantly reduced (∼50%) the PP1-PNUTS interaction (Fig. 3C and S6B), supporting the importance of the hydrophobic interface with the conserved Val and Trp moieties of the LtPNUTS RVxF motif. We suggest that the VxW motif in LtPNUTS is the putative counterpart of a Vx(F/W) motif that comprises a key part of the PP1 phosphatase-binding site identified in several other PP1 regulatory subunits, including hPNUTS, where the VxW motif binds to a hydrophobic pocket of the phosphatase remote from the phosphatase active site (17) (Fig. S5B).

**Figure 3.**
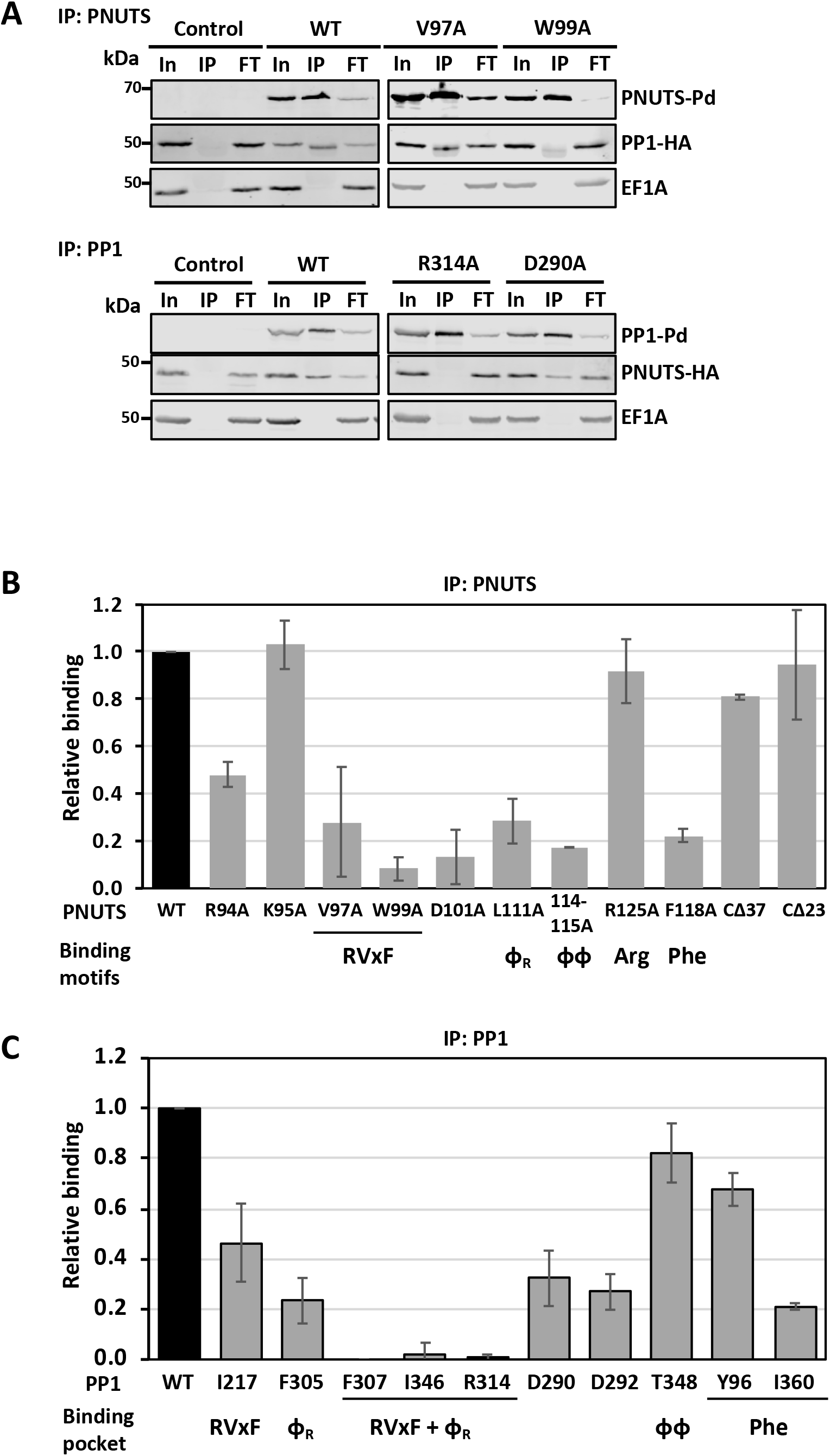
LtPNUTS binds LtPP1-8e using an extended RVxF-. Φ**_R_-**[[**-Phe motif.** A, co-immunoprecipitation assay of PP1-8e binding to PNUTS and their derivatives. PNUTS IP; PP1-8e was endogenously tagged with HA tag, and wild type or indicated PNUTS mutants with Pd tag were over-expressed from the pSNSAP1 vector. Cell extracts from the indicated cell lines were purified by anti-protein A affinity resin and analyzed by western blot with anti-protein A and anti-HA. In; input (equivalent to the amount of protein added to the IP reaction mixture), IP; 100% of the precipitated immunocomplexes, FT; flow through or non-bound supernatant. EF1α provides a loading control and negative control for the IP. PP1 IP; PNUTS was endogenously tagged with HA tag, and wild type or indicated PP1-8e mutants with Pd tag were over-expressed from the pSNSAP1 vector. The levels of PNUTS pulled down in the PP1 IP were assessed by western blot as described above. Additional PNUTS and PP1-8e mutations analyzed by Co-IP are shown in Fig. S6. B and C, the relative binding (%IP) between PNUTS and PP1-8e (WT and variants) determined by the ratio of the band intensity of IP to that of In. B, PNUTS-Pd IP. The bar graph represents the mean ± SD from three independent experiments, with the % IP of PP1 using WT PNUTS set to 1. The PNUTS binding motif that corresponds to the residue tested, according to the model in Fig. 2, is indicated at the bottom of the graph. CΔ23 and CΔ37 refer to C-terminal truncations of PNUTS described in Fig. 4. C, PP1-8e-Pd IP. The % IP of PNUTS from the PP1 pull-down (WT and mutants) was determined as in B. The PNUTS binding pocket represented by each residue of PP1 is indicated below. F307, I346 and R314 are predicted to be key residues of both the RVxF and Φ_R_ binding pocket of PP1-8e.

A short 22-aa peptide from PNUTS that contains the RVxF motif is able to disrupt the PP1-PNUTS association, while the identical peptide with V97A and W99A substitutions is not (55), further confirming the importance of the RVxF motif in the LtPNUTS:PP1-8e complex. However, wild-type RVxF peptide did not elute all of the PP1 from PNUTS suggesting there may be additional interaction sites that stabilize the PNUTS–PP1 complex. PP1 phosphatase-regulatory proteins often have at least one, and often several, basic amino acids preceding the Vx(F/W) motif (69). It has been suggested that this basic region may interact with a negatively charged patch near the RVxF-binding pocket of PP1. In the case of hPNUTS, there is a run of five basic amino acids upstream of VxW (Fig. 2A).

Two of which engage in salt bridges to acidic side chains of PP1 (Fig. S5G). Similar interactions are predicted for LtPNUTS ^94^RKR^96^, which are predicted to have electrostatic interaction with PP1-8e residues D290, E340 and D292, respectively (Fig. S5H). Alanine mutation of R94A_PNUTS_, or its predicted interacting residue D290_PP1_, leads to a ∼50% reduction in PP1-PNUTS association (Figs. 3, B and C and S6, A and B). Alanine mutation of K95_PNUTS_ did not affect PP1-PNUTS interaction (Figs. 3B and S6A). However, not all electrostatic interactions mediated by these basic residues contribute equally to the association, and sometimes, simultaneous alanine mutations of all the basic amino acid residues preceding the RVxF motif is required to affect PP1 binding, as observed for the fission yeast PNUTS (70). We were unable to generate the R96A_PNUTS_ mutant, but alanine mutation of its interacting residue D292A_PP1_ leads to 60% reduction in PP1-PNUTS interaction (Fig. 3C and S6B), supporting the importance of R96_PNUTS_. Acidic residues C-terminal to the RVxF motif are also present in other PP1 regulatory subunits and important for binding PP1. The AlphaFold model shows D101 of LtPNUTS engaging in salt bridge interactions to R314_PP1_ (Fig. S5H). Similar interaction is observed on E405 of hPNUTS (Fig. S5G). Consistent with the prediction, the D101A_PNUTS_ mutant showed roughly 80% decreased interaction with PP1-8e (Fig. 3B and S6A).

The LtPNUTS:PP1-8e model predicts H114_PNUTS_ and V115_PNUTS_ form the PNUTS ΦΦ motif, which binds the PP1 ΦΦ binding pocket (Fig. 2E). Like the RVxF interaction, the ΦΦ interaction is highly similar to those observed in other PP1 holoenzyme complexes (Fig. 2C). The LL motif usually consists of two hydrophobic residues of PIPs that are buried in a hydrophobic pocket on PP1, but can be degenerate, including sequences such as VS, VC, VK, IN, and HH (17). The LL motif of hPNUTS is represented by ^410^YF^411^ located on a short β strand that hydrogen bonds with β strand β14 of PP1, extending one of its two central β sheets (Fig. S5C). AlphaFold predicts a similar arrangement in the LtPNUTS-PP1 complex (Fig. 2E and S5C). The predicted LL motif of LtPNUTS, ^114^HV^115^, is located on a short β strand formed by ^112^VKHV^115^ that potentially H-bonds with PP1-8e’s β strand 14 and the LL hydrophobic pocket on LtPP1-8e includes residues N100, R104, E350 and T348 (Fig. 2E). To test the significance of the LL motif, we mutated LtPNUTS ^114^VH^115^ to alanine, and found that the mutation significantly weakens the PP1-PNUTS association (Fig. 3B and S6A). While the structure of the hPNUTS:PP1 complex does not indicate any specific interactions between the LL motif of hPNUTS and the LL hydrophobic pocket of PP1, we noticed a potential salt bridge interaction between H114_PNUTS_ and T348_PP1_ in our model (Fig. 2E). Alanine mutation of T348_PP1_, however, had minimal impact on PP1-PNUTS association (Fig. 3C and S6B). Presumably, the interaction does not occur or the alanine mutation of T348 alone is not sufficient to disrupt the stabilizing β sheet interactions provided by the remaining residues in the pocket (Fig. 2E). A distinctive feature of the predicted LtPNUTS-PP1-8e structure is the extended linker between the RVxF and ΦΦ motifs (Fig. 2A). In LtPNUTS these two motifs are separated by 14 residues, and would represent the longest insert observed thus far for any PP1 regulator. In G_M_, these two motifs are separated by 10 residues (Fig. 2A). This “extended kink” is presumably stabilized by hydrophobic interactions made by the Φ_R_ motif, represented by L111_PNUTS_, with the hydrophobic Φ_R_ pocket adjacent to the RVxF binding pocket in PP1-8e (Fig 2B and 2D). As such, L111_PNUTS_ is stabilized by F305_PP1_, F307_PP1_, and R314_PP1_ components of the Φ_R_ pocket in PP1-8e (Fig. 2D). The contact mediated by the Φ_R_ motif (L407_hPNUTS_) is conserved in the hPP1:PNUTS structure (Fig. S5B).

Highlighting the importance of the L_R_ motif in the LtPNUTS:PP1 structure, L111A_PNUTS_ mutation significantly reduced (80%) the PNUTS-PP1 interactions (Figs. 3B and S6A). Furthermore, alanine mutation of F305_PP1_ decreased PP1- PNUTS association by 80%, and single alanine mutations of residues lining both the RVxF and Φ_R_ pockets of PP1-8e, F307, I346 and R314, completely disrupts the LtPP1-PNUTS association (Figs. 3C and S6B).

In many cases, PP1 interactions can extend beyond the ΦΦ motif. For example, F413_hPNUTS_ is the Phe motif that binds in a deep pocket immediately adjacent to P298_PP1_ in the human complex (Fig. S5D). This pocket is also frequently used by other regulators to bind PP1. For example, Gm (F82_GM_)(68), spinophilin (T461_spino_)(67), and RepoMan/Ki67 (F404_RM_)(59) and RRP1B (F696_RRP1B_)(61) bind this same pocket. A conserved F118 is present in LtPNUTS and predicted to bind a pocket adjacent to P351_PP1_ in the LtPNUTS:PP1 model (Fig. 2F). P351_LtPP1_ occupies a similar position as P298_hPP1_ in the human PP1 pocket (Fig. S5D). Additionally, the model predicts that Y96, N100, R104, L349, P352 and I360 form the Phe binding pocket in LtPP1-8e (Fig. 2F). While the Y96A_PP1_ mutation had a minor effect, alanine mutation of F118_PNUTS_, or I360_PP1_ significantly reduced LtPP1-PNUTS associations (Fig. 3B, 3C, S6A and S6B), supporting a similar involvement of the Phe motif in the LtPNUTS-PP1 complex.

An additional potential LtPP1-8e interaction beyond the ΦΦ motif is R125_PNUTS_ (Fig. 2A). In hPNUTS R420 is involved in hydrophobic and electrostatic interactions with PP1, representing the so-called Arg motif (Fig. S5E).

R420_hPNUTS_ is buried in a hydrophobic pocket formed by L296, P298, and P270 of hPP1. Additionally, E419_hPNUTS_ and R420_hPNUTS_ form bidentate salt bridges with R74_PP1_ and D71_PP1_, respectively (Fig. S5E). However, this interaction is not predicted by AlphaFold in the LtPNUTS:PP1-8e complex. Rather, an alpha helix (residues 354-362) within the C-terminal tail of PP1-8e occupies the PP1 hydrophobic pocket involved in R420 hPNUTS binding (Fig. S5F).

Furthermore, while the Arg motif is presumably conserved on LtPNUTS as R125, the polar S124_PNUTS_ replaces the negatively charged E419_PNUTS_ in hPNUTS. Moreover, the interacting charged residue in hPP1, R74_PP1_, is replaced by N100 in LtPP1-8e (Figure S7). The replacement of charged residues with polar residues may prevent the formation of a bidentate salt bridge important for Arg motif binding. This concept along with the blocking of the Arg pocket by the C-terminal tail of LtPP1-8e could explain the divergence of the LtPNUTS-PP1 binding structure from hPNUTS at this region. R125 on LtPNUTS is therefore not predicted to bind to PP1-8e, and alanine mutation of R125 in LtPNUTS had no effect on PP1-PNUTS association (Figs. 3B and S6A). Alanine mutation of R420_hPNUTS_, however, does not affect PP1-PNUTS association in human cells (17), although the crystal structure indicates the importance of the Arg motif. Therefore, while the AlphaFold model clearly rules out the interaction, our Co-IP results do not completely exclude the possibility that R125_PNUTS_ mediates interaction with PP1 in *L. tarentolae* cells.

Taken together, the predicted structure and mutagenesis analyses establishes that LtPNUTS, like a majority of PP1-specific regulators, binds LtPP1-8e, in part, using a general RVxF and L_R_-ΦΦ-F SLiMs. We noticed that a majority of the mutant PNUTS proteins tested here are over-expressed at a lower protein level than WT PNUTS protein (Fig. S9A). This is not observed for PP1 mutants over-expressed from the same plasmid (Fig. S9B), suggesting that PNUTS protein level is sensitive to mutations. However, reduction in the level of over-expressed PNUTS does not necessarily lead to reduction in PP1 binding in the co-IP. For example, R125A_PNUTS_ is one of the lowest expressed PNUTS mutants (Fig. S9A) but showed comparable PP1 association as WT PNUTS (Fig. 3B). Potentially, the reduced expression of the mutant PNUTS protein is sufficient for saturation binding of available PP1, allowing co-IP of PP1 to the same extent as WT PNUTS. Furthermore, confirmation of the PNUTS-PP1 interface based on mutation of PNUTS residues is supported by mutation analysis of the corresponding binding pocket on PP1 where expression levels are not affected by mutagenesis. Therefore, the reduced protein expression level of certain mutant PNUTS protein does not affect our overall conclusions regarding the RVxF-L_R_-ΦΦ-F motifs.

### PP1-8e isoform specific residues are involved in PNUTS binding

The mode of PP1-PNUTS interaction described above, via the established RVxF-L_R_-ΦΦ-F motif, is typical for a scaffolding function of regulatory proteins but likely does not affect selectivity toward PP1 isoforms. In fact, a majority of the PP1 residues characterize above as involved in PNUTS-PP1 binding are not restricted to the PP1-8e isotype (Fig. S7), and thus, fail to explain the marked preference of PNUTS for PP1-8e. Therefore, we explored structural features present in LtPP1-8 that would confer specificity to LtPNUTS. As mentioned above, hPP1 isoforms share a high sequence identity with differences mainly limited to their extremities and some PIPs take advantage of these differences to interact selectively with specific PP1 isoforms. As we recently noted (55), an interesting characteristic of the PP1 isotypes in Leishmania is the diversity of their N- and C-terminal tails and the insertion of several short sequence elements specifically within the catalytic subunit of PP1-8e (Fig. S2). To test the contribution of each of these unique PP1-8e characteristics to LtPP1-PNUTS association, we performed deletion and alanine mutagenesis (constructs used in this study are illustrated in Fig. 4A). As shown in Fig. S8B, all PP1-8e mutants are over-expressed in leishmania cells to similar levels as WT PP1-8e. Upstream of the highly conserved catalytic domain, LtPP1-8e has a 32 amino N-terminal extension (Fig. S2). Deleting the PP1-8e N-terminal region completely abolished PP1-PNUTS association by co-IP (Figs. 4B and S9), suggesting that residues 1-32 of LtPP1-8e are essential for this interaction.

**Figure 4.**
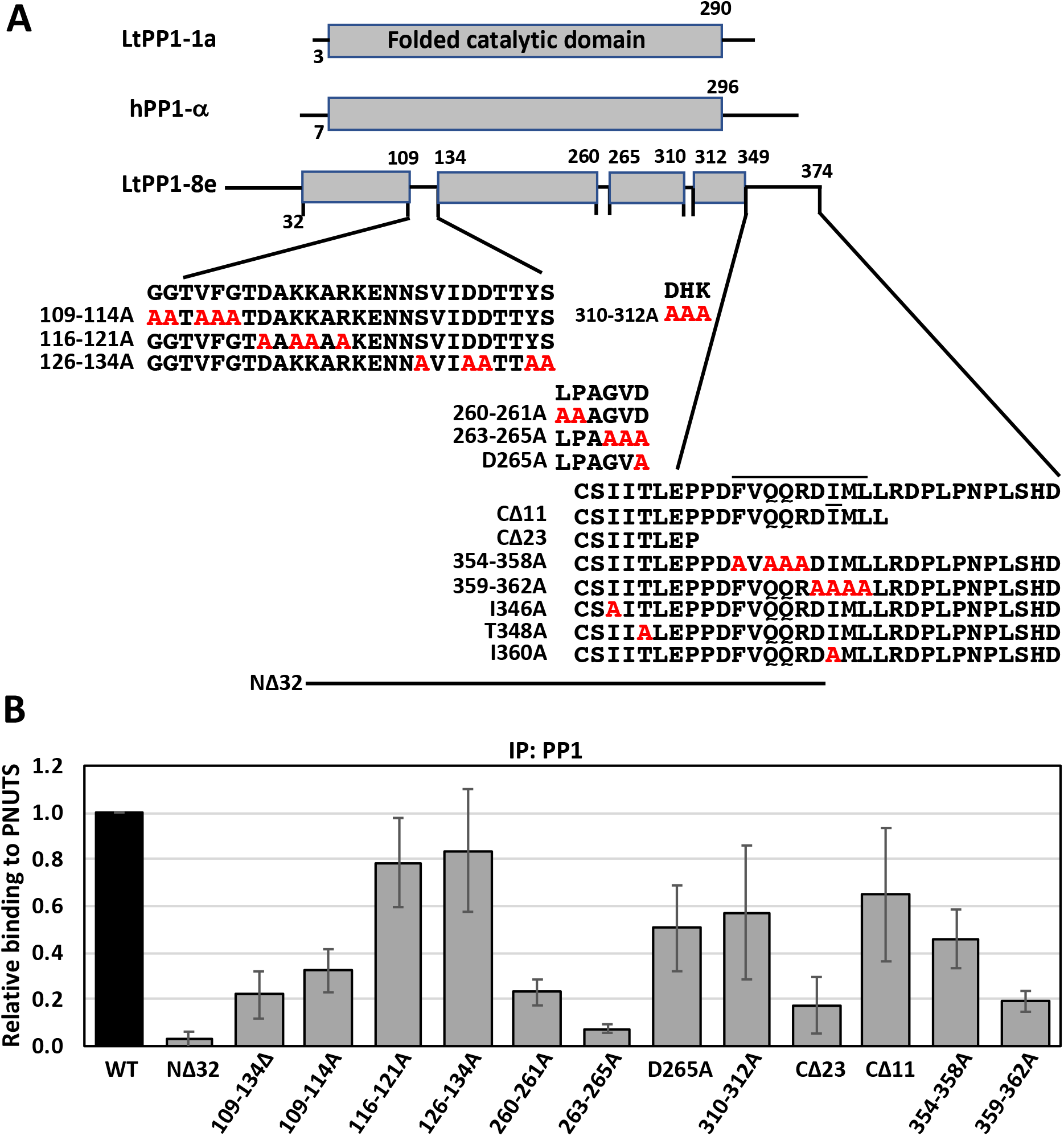
Non-canonical sites on PP1-8e are essential for PNUTS binding. A, PP1-8e constructs. The conserved PP1 catalytic domain is shown as grey boxes. Isoform differences between LtPP1-8e and LtPP1a and hPP1 are indicated by the lines within the catalytic domain and at the N- and C-terminus of LtPP1-8e. Sequence is provided for all these regions in LtPP1-8e, except the N-terminus, and residues subjected to alanine mutagenesis (red) or deletion are indicated. Residues in the predicted α-helix at the C-terminus are indicated by the line above the sequence. B, The % IP of PNUTS from the PP1 pull-down (WT and indicated variants) was determined as described in Fig. 3C. The bar graph represents the mean ± SD from three independent experiments, with the %IP from wild type PP1 set to 1. See Fig. S10.

Sequence differences between mammalian PP1-α and PP1-γ C-terminal ∼25-amino-acid tails are implicated in isoform specific binding by ASPP2 (60) and MYPT1(56). Similarly, Lt PP1-8e has a unique extended C-tail of ∼25 amino acids (Fig. S2) that includes two residues (P352 and I360) that we have demonstrated above as important for PNUTS-PP1 binding, potentially via stabilization of the Phe motif (Fig. 2F). Deleting the PP1-8e C-terminal tail significantly impacted PNUTS binding *in vivo* (Fig. 4B). While deletion of the 23 amino acid C-terminal extension of PP1-8e (CΔ23) leads to ∼80% loss in PNUTS binding, deletion of the final 11 amino acids (CΔ11) resulted in ∼30% reduction in PNUTS binding (Figs. 4B and S9). The 12 amino acid of the C-term region between these two deletions includes a predicted 9 amino acid alpha-helix (354–362) rich in charged or polar residues (Fig. S2), potentially involved in electrostatic interactions with PNUTS. To test this idea, we did alanine scanning mutagenesis of two regions within this C-terminal helical region. Alanine substitution of four residues within first half of this helix in PP1-8e (354-358A) resulted in 50% reduction in PNUTS binding and the 359-362A mutation of PP1-8e resulted in 80% reduction in PNUTS binding, similar to what we observed in the 23 amino acid deletion (CΔ23) (Fig. 4B). The I360A_PP1_ mutant led to a similar 80% reduction in LtPNUTS-PP1-8e associations, indicating I360 is a key residue within this C-term 359-362 helical region. Thus, the unique C-terminal tail of PP1-8e, in particular residues 359-362, and the first 32 amino acids at the N-terminus are needed for PNUTS binding.

According to the LtPNUTS:PP1-8e model, while the N-terminus of PP1-8e is unstructured and thus, difficult to understand how it is involved in isoform selective binding to PNUTS, the C-terminus appears to provide additional stabilization to the Phe binding pocket. PP1-8e, and other LtPP1 isotypes, have an Phe binding pocket similar to the human PP1:PNUTS complex (Fig. 2F and Fig. S5D and S7). However, the unique C-terminus of PP1-8e provides additional residues (including P352 and I360) that may contribute to the Phe binding pocket. To examine this idea further we determined the AlphaFold model for the LtPNUTS:PP1-1a complex (Fig. S10A). LtPP1-1a appears to have a majority of the conserved residues for the RVxF, L_R_, LL and F motif binding pockets as the human PNUTS:PP1 complex and the predicted LtPNUTS:PP1-8e model (Figs. S7 and S10). However, LtPP1-1a lacks the extended C-terminus present in LtPP1-8e (Figs. S2 and S10C) and, interestingly, is predicted to associate with LtPNUTS with the RVxF-L_R_-LL motifs, but not the Phe motif (Fig. S10)

An additional characteristic of PP1-8e is the insertion of three unique sequence motifs within the catalytic domain; a 26 amino acid insertion (residues 109-134) near the N-terminus and two smaller (^260^LPAGVD^265^ and ^310^DHK^312^) insertions near the C-terminus (Fig. S2). AlphaFold modeling shows the insertions are presented on the surface of PP1-8e at novel sites compared with the human PP1 structure and the LtPP1-1 isoform (Fig. S3, E and F). To test the significance of these regions, we performed deletion and alanine mutagenesis. Deletion of the 26 amino acid insertion in PP1-8e (109-134Δ) results in severely reduced ability (80%) to associate with PNUTS (Fig. 4B). The 26 amino acid region is rich in charged and polar residues that are conserved among Leishmania PP1-8e homologs, potentially involved in electrostatic interactions with PNUTS. To test this idea, we did alanine mutagenesis in three regions of the 26 amino acid insertion: GGVFG (109-114A), DKKR (116-121A) and SDDYS (126-134A) (Fig. 4A).

While the 116-121A and 126-134A mutations had little effect on PNUTS binding, mutation of five residues in 109-114A resulted in 80% reduction in PNUTS binding, similar to the effect of deleting the entire 26 amino insert (Fig. 4B).

Similar alanine mutagenesis was performed for the two smaller PP1 insertions: ^260^LPGVD^265^ and ^310^DHK^312^ (Fig. 4B). The results show that while alanine mutagenesis of ^310^DHK^312^ leads to a small decrease (∼20%) in PP1-PNUTS association, alanine mutagenesis of ^260^LP^261^ abolishes roughly 80% of PP1-PNUTS interaction (Figs. 4B and S10). Alanine substitution of the remaining three residues of the ^260^LPGVD^265^ (263-265A) led to approximately 90% reduction in PNUTS binding, and D265A_PP1_ mutation only had a moderate effect on PP1-PNUTS interaction (Figs. 4B and S10). Thus, unique sequences within the catalytic domain of PP1-8e, in particular residues ^109^GGTVFG^114^ and ^260^LPAGV^264^, are needed for PNUTS interaction. Taken together these results suggest that LtPNUTS can discriminate between different PP1 isoforms based on the PP1 N- and C-terminus and unique sequence motifs within the catalytic domain.

As such, these regions might underlie the mechanism by which LtPNUTS shows preferential binding to PP1-8e.

### PNUTS as a scaffold for the PJW/PP1 complex

hPNUTS is a scaffolding protein in the human PTW/PP1 complex, binding Tox4 and Wdr82 with its N- and C-terminus regions, respectively, and PP1 via the centrally located RVxF motif (21). The hPNUTS is a 114 kDa protein (940 amino acids) with multiple identified protein domains (13). LtPNUTS, which lacks identifiable protein domains or motifs apart from the conserved PP1-interacting RVxF motif discussed above, is much smaller at 28.6 kDa, consisting of 264 amino acids. To test if LtPNUTS similarly serves as a scaffolding protein and binds to Wdr82 and JBP3 with distinct domains we over-expressed Pd-tagged PNUTS protein with various of N- and C-terminal truncations and studied the interaction between PNUTS truncations and endogenously HA-tagged JBP3/Wdr82 using Co-IP (Fig. 5). We find that full length PNUTS allows significant co-IP of both JBP3 and Wdr82 (Figs. 5C and S11), consistent with our previous studies of the PJW/PP1 complex in Leishmania and *T. brucei* (*53*). Confirming that the RVXF motif and PP1 binding are not required for JBP3 and Wdr82 association with PNUTS, mutation of the PP1 binding RVxF motif (RACA mutant) has little to no effect on Wdr82 or JBP3 binding to PNUTS (Figs. 5C and S11). Interestingly, PNUTS proteins with three different N-terminal truncations (NΔ27, NΔ47, and NΔ75) are expressed at significantly lower levels than the full-length PNUTS control, and the major species run at lower molecular weights than expected on SDS-PAGE gel (Fig. 5B and S8C). As an intrinsically disordered protein, hPNUTS is known to not run to the expected size on the SDS-PAGE gel (9) and we have characterized the altered mobility of TbPNUTS (53). Potentially, the deletion of an N-terminal sequence accentuates the disordered nature and altered mobility of the truncated LtPNUTS polypeptide. In this case, the major species represents the indicated truncated PNUTS protein. Alternatively, N-terminal deletions lead to PNUTS protein instability and further protein cleavage. While it is difficult to obtain accurate measurement of binding with such low protein expression in the parasite, it seems that PNUTS with varying lengths of N-terminal truncations still immunoprecipitated a significant level of Wdr82 or JBP3 compared to the negative control, although not to the same extent as WT PNUTS (as shown in Fig. S11).This is best represented by the NΔ75 PNUTS, with the highest level of expression among the N-terminally truncated PNUTS proteins (Fig. 5B). This would suggest the N-terminus of PNUTS is not essential for JBP3/Wdr82 binding. We noticed that similar N-terminal truncations of the PNUTS homolog in *T. brucei* does not result in decreased levels of expression (Fig. S12, A and B), allowing further studies of Wdr82/JBP3 association. To do this we tagged JBP3 and Wdr82 in *T. brucei* with HA and Myc tags respectively, and exogenously expressed protein A-tagged PNUTS via a Tet-inducible promoter. Supporting the LtPNUTS analysis, 72 aa deletion from the N-terminus (NΔ72) tested in TbPNUTS had little to no effect on Wdr82/JBP3 binding (Fig. S12C). In contrast, while all C-terminal truncations of LtPNUTS are expressed at levels similar to full-length in both HA tagged Wdr82 and JBP3 cell lines (Fig. 5B and S8C), even the smallest 23 amino acid deletion (CΔ23) had a negative effect on both Wdr and JBP3 binding to LtPNUTS (Figs. 5C and S11). While the 23 amino acid deletion led to complete loss of JBP3 binding, a small level of Wdr82 association remained that is subsequently lost upon further deletions of the C-term end (CΔ37 and CΔ66) (Figs. 5C and S11). Similar to LtPNUTS, C-terminal deletion of TbPNUTS (CΔ82) results in complete loss of JBP3/Wdr82 association (Fig. S12C). CΔ23 and CΔ37 PNUTS had a minor but insignificant effect on LtPNUTS-PP1-8e association (Figs. 3B and S6A), suggesting that PP1 binding into the complex is not dependent on Wdr82 or JBP3 interaction. Taken together the data suggests the C-terminus of LtPNUTS (and TbPNUTS) is required for binding both Wdr82 and JBP3 and that binding is independent of PP1 binding at the central RVxF motif.

**Figure 5.**
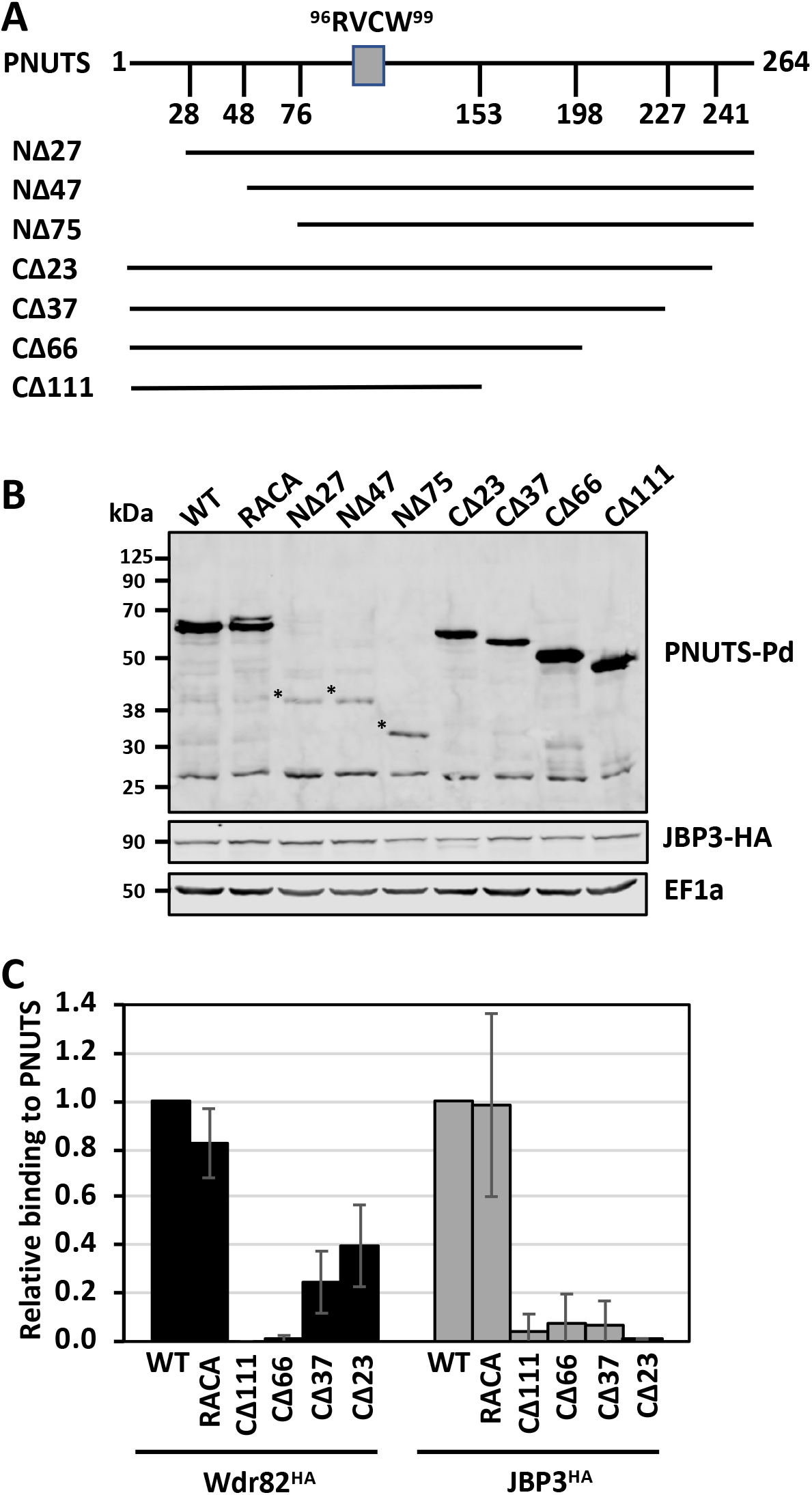
JBP3 and Wdr82 bind to the C-terminus of LtPNUTS. A, schematic diagram of PNUTS depicting the PP1- specific RVxF SLiM (RVCW). Constructs used in this study are illustrated. Underlined residues at the C-terminus indicate residues subjected to alanine mutagenesis. B, western blot showing the protein expression of Pd-tagged PNUTS (WT and truncation mutants) in JBP3-HA tagged *L. tarentolae* cells. Dots indicate the proposed products representing the indicated N-terminal truncations. Anti-EF1A western blot is shown as a loading control. C, analysis of JBP3/Wdr82 binding to PNUTS by Co-IP. PNUTS truncations (C) or mutants (D) were tested for interaction with HA-tagged Wdr82 or JBP3 by Co-IP analysis. %IP of WT Wdr82 and JBP3 by PNUTS were set to one and relative %IP of the indicated mutants was determined as described in Fig. 3. The bar graph represents the mean ± SD from three independent experiments. See Fig. S9.

Thus far, we have been unable to produce soluble recombinant protein in *E. coli* to study the PJW/PP1 complex formation in vitro. Therefore, to further test the scaffold function of PNUTS in the complex and clarify its binding relationship with Wdr82 and JBP3, we utilized the RNAi system in *T. brucei*. This system would allow us to characterize, for example, the effect of PNUTS knock-down on the interaction between Wdr82 and JBP3 by Co-IP. Therefore, we tagged JBP3 and Wdr82 with HA and Myc, respectively, in the PNUTS RNAi cell line. We find that knock-down of TbPNUTS leads to decreased protein levels of both Wdr82 and JBP3 (Fig. 6A). JBP3 is particularly sensitive to PNUTS knockdown, with the majority (>90%) of JBP3 being lost within 24h of PNUTS RNAi induction. On the other hand, Wdr82 is less affected with 50% reduction in protein level within 24h, with levels decreasing to ∼75% reduction upon 72 hr post induction. While this effect does prevent the analysis of JBP3/Wdr82 interactions by Co-IP, it is consistent with PNUTS knockdown in HEK293 cells which leads to loss of both Tox4 and Wdr82 (21), and further supports a scaffold function for LtPNUTS. Interestingly, knockdown of Wdr82 by RNAi similarly leads to a significant reduction in HA-tagged JBP3 protein level, but does not affect Myc-tagged PNUTS protein level (Fig. 6B). On the other hand, ablation of JBP3 by RNAi does not lead to any change in PTP-PNUTS or Myc-Wdr82 protein levels (Fig. 6C).

**Figure 6.**
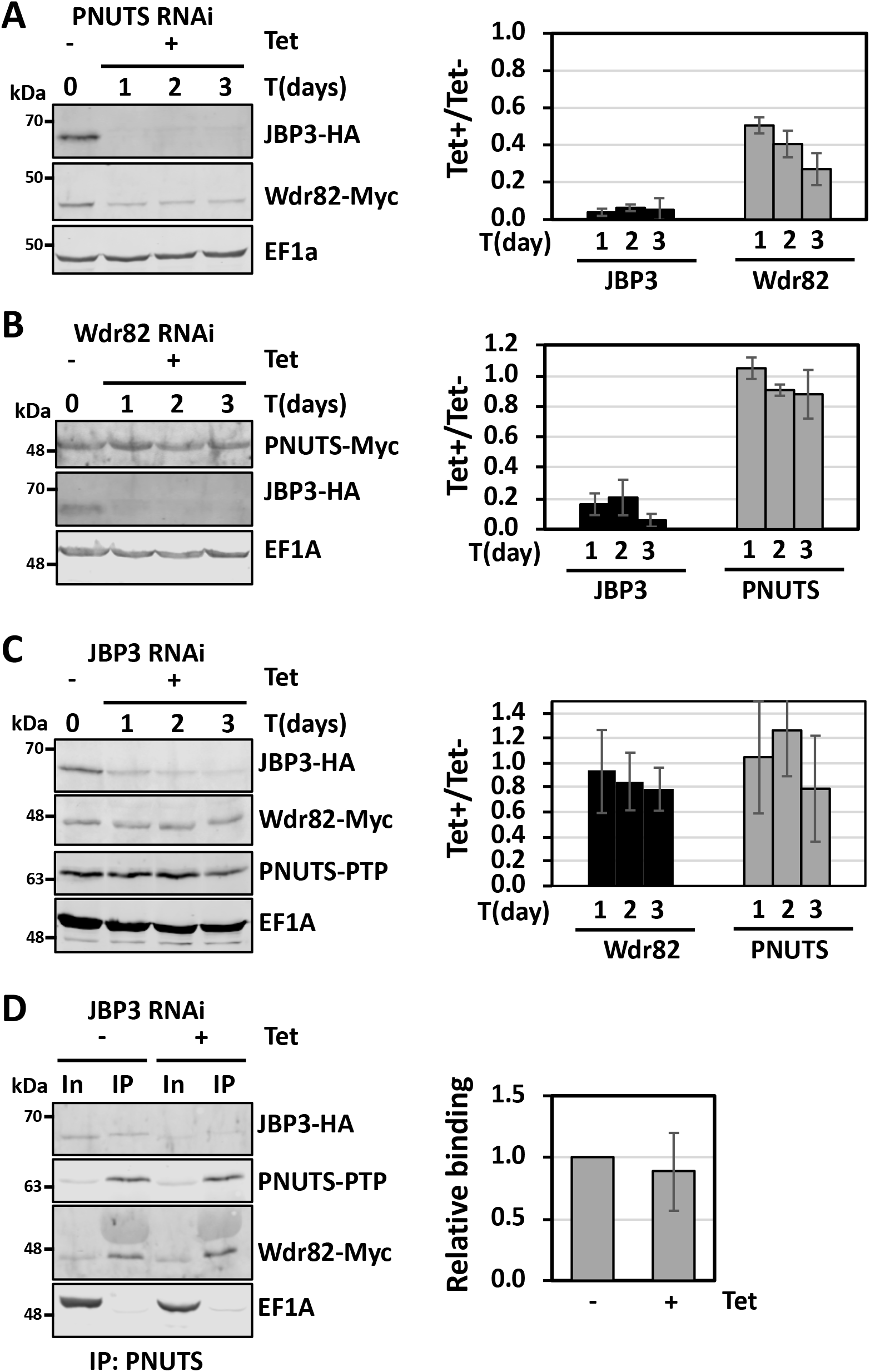
TbPNUTS functions as a scaffold factor. RNAi knockdown of the *T. brucei* PJW complex components. Endogenous loci of the indicated genes were tagged with HA, PTP, or Myc tags. Cells were then transfected with the indicated RNAi construct and knockdown of PNUTS (A), Wdr82 (B) or JBP3 (C) was induced by tetracycline (Tet) addition. Cell lysates were collected at the indicated time points and analyzed by western blot with anti-protein A, anti-HA or anti-Myc. Anti-EF1a serves as a loading control. Bands were quantified by densitometry. The bar graphs on the right represent the mean ± SD from 3 independent experiments for the indicated protein level relative to protein level prior to the induction of RNAi. D, effect of JBP3 knockdown on PNUTS-Wdr82 binding. JBP3 RNAi was induced for 48 hrs, and PNUTS-PTP was purified from cell extracts by anti-protein A affinity resin and analyzed by western blot. The %IP of Wdr82 by PNUTS IP with or without JBP3 RNAi induction was determined as described in Fig. 3. The bar graph on the right represents the mean ± SD from three independent experiments, with the %IP from the uninduced cells set to 1.

PNUTS and Wdr82 association was analyzed by anti-protein A Co-IP with or without JBP3 RNAi induction. The result shows that JBP3 knockdown does not affect PNUTS-Wdr82 Co-IP (Fig. 6D), indicating that Wdr82 binds to PNUTS independently of JBP3. The results collectively further suggest that JBP3 associates into the complex via binding to Wdr82, and that complex integrity is essential to Wdr82 and JBP3 protein stability.

In setting up the PP1-PNUTS Co-IP analysis and over-expressing LtPNUTS from a plasmid in cells expressing a tagged version of PP1-8e from the endogenous locus, we noticed that transfection with the PNUTS expressing plasmid led to a ∼50% decrease in PP1-8e abundance (Fig. 7, A and B). PNUTS over-expression has no effect on PP1-7 protein level, indicating an isotype specific effect. Interestingly, the effect of PNUTS over-expression on PP1-8e levels is not dependent on PP1 binding, since this occurs even upon over-expression of the PNUTS defective for PP1 association, such as RACA PNUTS (Fig. 7A), L111A_PNUTS_, F118A_PNUTS_, or R125A_PNUTS_ (Fig. S13A). However, over- expression of C-terminal truncated versions of PNUTS (CΔ23 or CΔ37) did not lead to reduced PP1 protein level to the same extent as other tested PNUTS mutants, indicating that the effect is dependent on the ability of PNUTS to bind Wdr82 and/or JBP3. Furthermore, while PNUTS over-expression had no effect on Wdr82 protein abundance, in a few clones it led to a shift in mobility of Wdr82 on the SDS-PAGE gel (Figs. 7C and S13B). Endogenously HA tagged Wdr82 has a predicted molecular weight of 48 kDa, and a majority of the protein runs slightly above the 50 kDa protein ladder marker with a minor lower molecular weight species sometimes visible just below the marker (Fig. 7C). We observed that WT and RACA PNUTS over-expression caused the population of Wdr82 to shift to the lower molecular weight species (Fig. 7C) in 2 out of 9 and 6 clones analyzed, respectively (Fig. S13B). The finding that expression of WT LtPNUTS and the RACA mutants had similar effects on the altered mobility of Wdr82 *in vivo*, indicates that the effect is independent of the ability of PNUTS to bind PP1-8e. Treatment of cell lysates with or without calf intestinal phosphatase and conditions we have demonstrated to dephosphorylate RNA Pol II (55), had no effect on Wdr82 gel mobility (data not shown), excluding the possibility that the observed shift in Wdr82 is due to changes in phosphorylation status. The AlphaFold predicted Wdr82 structure indicates the N-terminus (1–27) of Wdr82 has low prediction confidence, followed by potentially solvent exposed ^34^FYTGIN^39^ sequence susceptible for cleavage by chymotrypsin and thermolysin (Fig. S14A), suggesting a disordered N-terminus region prone to proteolytic cleavage.

**Figure 7.**
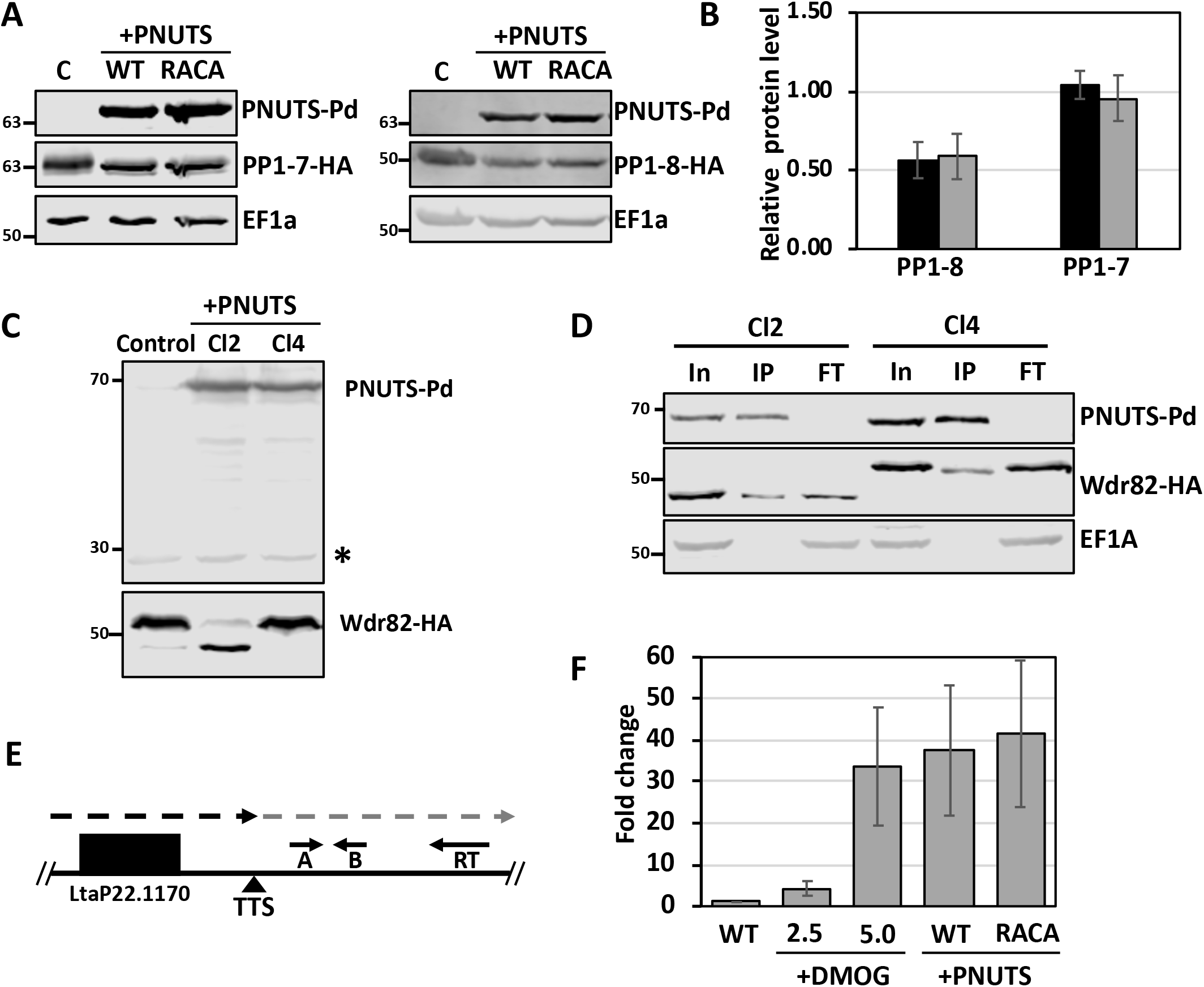
LtPNUTS overexpression alters PP1 and Wdr82 stability and transcription termination. A-D, effect of PNUTS overexpression on PP1 and Wdr82. A, PP1-1 or PP1-8 was tagged with HA-tag at its endogenous loci and either WT or RACA mutant PNUTS protein was exogenously overexpressed. Cell lysates were analyzed by western blot with anti-protein, anti-HA and anti-EF1a. Anti-EF1A serves as a loading control. PP1 tagged control cell lines not transfected with the PNUTS expression plasmid are indicated by the C for control. B, HA-tagged PP1-1 and PP1-8 band intensities were quantified by densitometry. The bar graph represents the mean ± SD of PP1-1 or PP1-8 protein level relative to control cells with no overexpression of PNUTS (WT, black bar; RACA mutant, grey bar). C, Wdr82 was tagged at its endogenous loci with HA tag with or without WT PNUTS overexpression and cell lines were analyzed by western blot with anti-protein A, and anti-HA. A non-specific product recognized by the anti-protein A antibody is indicated by an asterisk and serves as a loading control. Shown here are results from two clones (Cl2 and Cl4). See Fig. S13B for results from multiple clones. D, cell extracts from the cell lines in C were purified by anti-protein A affinity resin and analyzed by western blot with anti-protein A, anti-HA and anti-EF1a. E and F, effect of PNUTS overexpression on Pol II transcription termination. E, diagram of the termination site at the end of a polycistronic gene array on chromosome 22 illustrating the strand-specific RT-qPCR analysis of readthrough defects. The dashed line indicates the readthrough transcripts past the transcription termination site (TTS) that accumulate following a defect in Pol II termination. The location of primers for RT (RT) and qPCR (A and B) are indicated by the small arrows. F, RT- PCR analysis for the readthrough transcripts. Strand-specific cDNAs were synthesized from RNAs extracted from cells treated with the indicated concentrations of DMOG, or from cells with either WT or RACA mutant PNUTS overexpression using primer RT. Fold change of the readthrough transcripts relative to the WT ± SD is based on qPCR analysis with primer A and B, normalized to tubulin RNA.

Preliminary MS analyses to confirm the processing of the lower MW form of Wdr82 have been inconclusive. Regardless of the explanation for the different species of Wdr82 generated by LtPNUTS overexpression, both species bind LtPNUTS to similar degree. Co-IP studies show that the full-length and truncated Wdr82 species IP similar amounts of PNUTS (Fig. 7, C and D), indicating the potential cleavage of the Wdr82 N-terminus does not affect PNUTS association.

While it is unclear why LtPNUTS overexpression results in these effects on Wdr82 and PP1-8e, the data further support a scaffold function for PNUTS in the PJW/PP1 complex. Furthermore, as predicted based on previous studies of PJW/PP1 complex function *in vivo* in Leishmania and *T. brucei*, these defects correlate with defects in Pol II transcription termination (Fig. 7, E and F). Strand-specific RT-qPCR shows that compared to the parental cells (WT), cells that over-express WT or RACA PNUTS accumulated nascent transcripts downstream of the analyzed transcription termination site (Fig. 7, E and F). As a positive control, cells treated with DMOG, a drug that inhibits base J synthesis and induces transcription termination defects in Leishmania (49), also accumulated readthrough transcripts. The effect of PNUTS overexpression and corresponding decreased levels of PP1-8e on RNA Pol II termination seen here is consistent with the recently characterized role of PP1-8e in Pol II phosphorylation and transcription termination in *Leishmania* (*55*). To address the impact of Wdr82 cleavage that occurs following over- expression of PNUTS to the termination defects measured here, we repeated the analysis using HA-tagged Wdr82 cells and examined the degree of readthrough in cells with or without Wdr82 cleavage (Fig. S13, C and D). Compared to WT cells, C-terminal tagging of Wdr82 leads to increased readthrough transcripts (Fig. S13D), possibly indicating an impaired function for Wdr82-HA, similar to what we observed for C-terminally tagged PP1-8e in *L. major* (55).

However, we see no difference in the degree of readthrough transcription stimulated by PNUTS overexpression in cells that resulted in Wdr82 cleavage or not (Fig. S13C). Therefore, altered processing of Wdr82 in the PNUTS expressing cell lines had no additional negative effect on Pol II transcription termination. Taken together, similar to the termination defects measured in the Leishmania PP1-8e KO (55), alterations in PJW/PP1 complex formation and levels of PP1-8e following PNUTS over-expression lead to defects in Pol II transcription termination.

## Discussion

PP1 protein phosphatase has little intrinsic substrate specificity (3), allowing it to dephosphorylate diverse substrates and be involved in a wide range of biological activities. Its phosphatase activity is regulated *in vivo* in a spatiotemporal manner by interacting with PIPs (7). As such PIPs are essential regulators of PP1 substrate specificity and cellular localization. PIPs share little sequence or overall structural identity but use short SLiMS (5-8 amino acids long) that are combined within an unstructured domain to render PIPs high affinity to PP1 (8). According to this PP1 binding code (8), the unique combination of PP1 binding motifs (SLiMS) allows PIPs to interact with PP1 in a highly specific manner. PNUTS-PP1 complex involved in regulating transcription termination is conserved from mammalian to yeast cells (21, 71, 72), and recent studies indicate that the binary interaction and function also exists in trypanosomatids (53, 54), a group of early diverging eukaryotes. Purification of the PNUTS complex from *L. tarentolae* identified a specific interaction with the PP1-8e isoform among the eight encoded in the Leishmania genome (53, 54) suggesting that PNUTS selectively targets PP1-8e to the complex. However, the isoform selectivity of PP1 targeting in intact parasites had not been established. Here we show that PNUTS selectively targets PP1-8e to the complex and targeting requires both the non-isoform selective canonical PP1-binding motif and additional domains located throughout the PP1-8e sequence. Previous studies have shown that LtPNUTS is a highly disordered protein, and mutation of its putative RVxF motif disrupts its interaction with LtPP1-8e, indicating its importance in PP1-PNUTS interaction (53). In the current study, we used AlphaFold to predict the LtPNUTS:PP1-8e holoenzyme complex, and identified additional SLiMs beyond the canonical RVxF motif that are typically difficult to recognize based on sequence analysis alone, because they are short and highly degenerate. Our predicted LtPNUTS:PP1-8e holoenzyme complex and biochemical studies reveal that LtPNUTS binds PP1-8e using an extended RVxF-L_R_-LL-Phe motif used by several other PIPs including the human PNUTS:PP1 complex. Furthermore, our studies suggest additional interactions are involved that are atypical compared with any previously studied regulator. These include unique sequences at the ends and within the catalytic domain of PP1-8e that modulate isoform specific recruitment as well as increasing overall stability of the holoenzyme complex.

Mammalian PIPs are known to take advantage of the divergent extremities and sequence polymorphism of PP1 to interact preferentially to specific isoforms (PP1α, PP1β, PP1γ). PP1 isoform specificity can be achieved via one of two mechanisms recognized thus far: PIP recognition of the PP1 C-terminus or via binding of the regulator to a recently characterized β/γ specificity pocket within the PP1 catalytic domain. C-terminal regions of PP1 are typically unstructured and difficult to crystallize, making it difficult to understand how the termini are involved in isoform selectivity (66). However, the structure of ASPP:PP1 identified SH3 domain interactions with the PPII motif within an alpha helical turn at the PP1 C-terminus enabling isoform selectivity (60). Myosin phosphatase targeting subunit 1 (MYPT1) interacts with Y305 and Y307 in the C-terminus of hPP1β via two ankyrin repeats. The two residues are not conserved in hPP1 α and γ, thereby enabling MYPT1 to selectively bind to hPP1β (56). Selective PP1 isoform specificity by other mammalian PIPs is not due to the C-terminal tail but rather to one of the seven amino acids that differ in the catalytic domains of PP1 isoforms. For example, selective nucleolar localization of two of the mammalian PP1 isoforms (PP1β and PP1γ) is due to selective interactions of the ribosomal RNA processing 1 homolog b protein (RRP1B). Isoform specificity of RRP1B, similar to two other PP1γ-specific regulators (RepoMan and Ki-67), is due to a single amino acid difference in PP1 at position 20, which is an Arg residue in PP1γ/β and a Gln residue in PP1α (64). Structural studies indicated this Arg is part of a hydrophobic AF-binding pocket in PP1, located between the N terminus and the catalytic domain, that engages RRP1B residues ^714^AF^715^ downstream of the extended RVXF motif, allowing Arg20 to mediate the selectivity of PP1-γ-specific regulators (59, 61). The studies presented here indicate LtPNUTS takes advantage of the divergent extremities of PP1 and novel sequence inserts within the catalytic domain to interact preferentially to the PP1-8e isoform. However, predicted LtPNUTS:PP1 interactions are unique compared to mammalian PIPs discussed above. Sequence and structural modeling does not support a similar AF-binding pocket in PP1-8e and LtPNUTS seems to lack the presentation of similar residues to engage the AF pocket. Although no motif, such as the PPII motif, is apparent on the C-tail of LtPP1-8e, our model predicts an alpha helix within the unique C-term of PP1-8e that is essential for interaction with LtPNUTS and isoform selectivity.

Compared to the other seven LtPP1 isoforms, LtPP1-8e has a long and unique C-tail with residues 354-362 predicted to form an α-helix secondary structure, and the remaining 12 residues (363–374) unstructured. The model indicates that two residues within the C-term α-helix (P352 and I360) accommodate the Phe motif in LtPNUTS (F118_PNUTS_). Supporting this model, C-terminal deletion and alanine scanning mutagenesis of PP1-8e indicates the importance of residues 359-363 of this alpha-helical region in LtPNUTS-PP1-8e interactions. The strong negative effects of the F118A_PNUTS_ and I360A_PP1-8e_ mutants on complex formation further supports this idea. Therefore, although the residues that constitute the conserved RVxF, L_R_, LL and F motif binding sites are present in all LtPP1 homologs, the PP1-8e C-tail may provide a stabilizing force to the PNUTS Phe SliM and represent a significant component of isoform selectivity. While human PP1 isoforms have a short divergent N-terminus (∼6 amino acids), a role of the N-terminus in PIP binding or isoform selection has not been described in other systems. Deletion of the LtPP1-8e N-terminus (1-32) leads to a dramatic decrease in LtPP1-PNUTS association, indicating its significance, but how it mediates association with LtPNUTS is unclear. The low confidence of the AlphaFold structure for this region makes it difficult to understand how the N-terminus is involved in isoform selectivity. We also now identify unique inserts within the PP1-8e catalytic region (^260^LPAGVD^265^ and ^310^DHK^312^ and the 26 amino acid 109-134 motif) where deletion or alanine mutagenesis completely abolishes or significantly decreases PP1-PNUTS association. Mutagenesis analysis has indicated residues ^260^LPGV^264^ and ^109^GGTVFG^114^ as key residues within these inserted motifs, essential for PNUTS-PP1-8e complex formation. Overall, the results suggest that polymorphisms within the PP1-8e catalytic domain and N- and C-terminus are essential for PNUTS binding. As such, these regions might underlie the mechanism by which LtPNUTS shows preferential binding to PP1-8e. However, the position/orientation of the LtPP1-8 polymorphisms were, in some cases, predicted with low confidence by AlphaFold. Therefore, how they contribute to PNUTS association cannot be easily inferred. Interestingly, the LtPNUTS:PP1-8e model predicts two of these unique regions of PP1-8e (C-term and the 26 amino acid internal motif) to be in close proximity to region 116-121 of PNUTS that includes the Phe SLiM (F118) (Fig. S10C). The predicted role of the C-term forming an essential part of the Phe binding pocket is discussed above. Within the 26-amino acid insertion polymorphism in the PP1-8e catalytic domain, ^113^FG^114^ is predicted to be in close proximity of Y117_PNUTS_ (Fig. S10C). The importance of this region is supported by our co-IP studies where alanine mutagenesis of residues 109-114 of PP1-8e, in contrast to mutagenesis of the remaining part of this 26-amino acid insert, significantly affected PP1-PNUTS association (Fig. 4). The absence of both of these regions in LtPP1-1 could therefore explain the altered Phe motif interactions in the PNUTS:PP1-1 holoenzyme model (Fig. S10). Taken together, the data support two unique regions of the PP1-8e isotype making critical interactions with PNUTS Phe motif that may help explain the isotype specific stable association of the LtPNUTS:PP1- 8e complex.

As mentioned above, our model predicts LtPNUTS binds PP1-8e via RVxF-L_R_-LL-Phe motifs, similar to the human PNUTS:PP1 complex. hPNUTS, like most PIPs is able to bind all PP1 isoforms. Presumably, the additional contacts with PP1-8e specific sequences we describe here allow isoform specific binding of LtPNUTS. However, the conservation of residues involved in interactions with the extended RVxF motif in all 8 LtPP1 isoforms (Fig. S7) suggests, as described for mammalian isoform specific PIPs, some low level of PNUTS binding *in vivo* by the remaining PP1 isoforms. This characteristic would explain the ability of other PP1 isoforms to functionally compensate for the loss of PP1-8e in *Leishmania* (55). The PNUTS-PP1-8e complex has been shown to regulate transcription termination in *Leishmania* potentially through PP1-8e-mediated dephosphorylation of Pol II CTD (55). KO of PP1-8e in

*L. major* causes transcription termination defects, which can be rescued, albeit to a limited degree, by over-expression of PP1-1 or PP1-7(55). Both proteins have conserved residues constituting the RVxF motif-binding pocket, and are predicted to interact with PNUTS through a majority of the extended RVxF motif. However, they do lack the PP1-8e unique motifs we demonstrate as critical for the PNUTS-PP1-8e Co-IP, including the C-tail and the ^113^FG^114^ residues we predict essential for stable Phe SLiM binding and thus increase overall stability of the holoenzyme complex. Therefore, it is conceivable that while the enhanced affinity for PNUTS allows LtPP1-8e to outcompete other PP1 isotypes for PNUTS binding in the WT cell, in its absence the remaining PP1 isotypes can form unstable or transient interaction with PNUTS to partially compensate for the loss of LtPP1-8e. Similarly, the lack of these LtPP1-8e specific polymorphisms essential for the PNUTS-PP1 co-IP in all eight TbPP1 isoforms may explain the failure to identify a PP1 isoform associated with PNUTS in *T. brucei*. While TbPNUTS has a conserved RVxF motif, purification of PNUTS from *T. brucei* cells identified the Wdr82 and JBP3 homologs but no catalytic PP1 component (53). Knockdown of TbPNUTS, TbJBP3 or TbWdr82 led to defects in Pol II transcription termination (53). Thus, we predict that a similar mechanism of Pol II termination involving PP1 mediated Pol II dephosphorylation via the PJW/PP1 complex exists in

*T. brucei* as we characterized in *Leishmania*. The inability to demonstrate TbPNUTS-PP1 binding using co-IP suggests that the two proteins do not interact directly or interact in such a transient or weak manner that PP1 dissociates from the complex during affinity purification process. Although we cannot exclude the possibility that the *T. brucei* PNUTS complex functions without the association of the catalytic PP1 component, the presence of an RVXF motif in TbPNUTS and lack of the polymorphisms we demonstrate as critical for stable LtPNUTS-PP1-8e interactions in the Co-IP in all *T. brucei* PP1 isoforms support our model.

The PNUTS-PP1 complex in mammalian cells is found associated with structural factors Wdr82 and Tox4, forming the PTW/PP1 complex (21). hPNUTS is a large protein with multiple domains; including the RVxF motif (KSVTW) for PP1 binding, an N-terminal transcription factor II S-like (TFIIS-like) domain required for Tox4 binding, RGG motifs toward the C-terminus, and a single zinc-finger domain at the extreme C-terminus (13). hPNUTS serves as a scaffolding protein in the PTW/PP1 complex and its ablation in HEK293 cells causes a complete loss of Tox4 and a significant reduction in Wdr82 protein level (21). LtPNUTS, on the other hand, is much smaller with no recognizable domains other than the central RVxF motif and extended SLiMs identified here involved in PP1 binding. For the first time, we now describe that PNUTS performs similar scaffolding function in the PJW/PP1 complex in kinetoplastids, representing a key regulator of complex formation/stability. We show that ablation of TbPNUTS leads to a complete loss of JBP3, the counterpart of Tox4, and a 50% reduction in Wdr82 protein level. Moreover, over-expression of LtPNUTS leads to reduction in PP1-8e levels and processing of Wdr82 (discussed below). We demonstrate that JBP3 and Wdr82 bind the C-terminus of LtPNUTS and TbPNUTS, independent of PP1 binding. The LtPNUTS defective for PP1 binding (RACA) has no detectable loss of binding to Wdr82 or JBP3, and C-terminal mutants, unable to bind Wdr82/JBP3, bind PP1 with WT level of efficiency. While there is no apparent interaction between Wdr82/JBP3 and PP1, C-terminal deletions had a significant negative effect on both Wdr82 and JBP3 association, suggesting interdependence of PNUTS binding by Wdr82 and JBP3. Alternatively, structural alteration in PNUTS caused by C-terminal deletion could explain the negative effects on the binding of both factors. While we are not able to rule this out, the effect would have to be localized to the C-terminus as the deletions have no measurable effect on PP1 binding. Furthermore, the use of RNAi in *T. brucei* supports the interdependence of PNUTS binding by Wdr82 and JBP3, where primary interactions between PNUTS and Wdr82 regulate JBP3 binding. While the ablation of JBP3 has no effect on Wdr82 levels or interactions with TbPNUTS, Wdr82 ablation leads to specific decrease in JBP3.

Presumably, in *T. brucei*, the stability of JBP3 depends on interactions with Wdr82 (and PNUTS). Additional work is needed to fully elucidate specific interactions involved in PJW/PP1 complex formation. However, taken together, the results from the *in vivo* studies suggest that PNUTS is a scaffolding protein in the PJW/PP1 complex that mediates the independent binding of PP1 and Wdr82, and JBP3 association with the complex depends, at least partially, on interactions with Wdr82.

The effects of overexpression of LtPNUTS, and ablation of TbPNUTS, on the PJW/PP1 complex supports its key role as a scaffolding factor for the complex and indicates the concentration of PNUTS is finely tuned *in vivo* in kinetoplastids. Presumably, over-expression of PNUTS in Leishmania leads to stoichiometric imbalance that affects PJW/PP1 complex formation and stability of associated factors, including PP1-8e. LtPNUTS over-expression had no detectable effect on levels of PP1-7 isotype that is not associated with the PJW/PP1 complex. Interestingly, the specific decrease of LtPP1-8e protein level is not dependent on the ability of LtPNUTS to bind PP1, but on its ability to bind Wdr82/JBP3. Over-expression of C-terminally truncated LtPNUTS (CΔ23 and CΔ37) with significantly lower affinity to Wdr82/JBP3 did not lead to a loss of LtPP1-8 as seen following over-expression of WT PNUTS or PP1-8e binding mutants. These results suggest that the integrity of the PJW/PP1 complex is important for PP1-8e protein level. Excess LtPNUTS (regardless of its ability to bind PP1) could lead to decreased levels of Wdr82/JBP3 (or other unidentified co-factors) available to PNUTS-PP1 to form a stable functional complex. The shift in molecular weight of Wdr82 in a percentage of clones overexpressing PNUTS is currently unclear. We have addressed the possibility of a shift due to phosphorylation and proposed it represents proteolytic cleavage at the unstructured N-terminus. Further work is needed to understand the effect of LtPNUTS over-expression on Wdr82 processing. However, this effect is not linked to the ability of PNUTS to bind PP1. While it is unclear if this altered Wdr82 processing affects cellular function, it has no apparent consequence on the ability of Wdr82 to bind PNUTS.

Overall, the current study identified PP1-binding motifs on LtPNUTS and discovered novel sequences on PP1- 8e that could confer isoform selectivity, thereby enhancing our understanding of the PP1 binding code modulating the interaction between PP1 and PP1-interacting proteins. Moreover, our results indicate the conserved role of PNUTS as a scaffolding protein and that its protein level is critical for PJW/PP1 complex stability. The finding that PJW/PP1 complex defects associated with PNUTS over-expression led to readthrough transcription at Pol II termination sites provides additional support for the involvement of the complex in the mechanism of Pol II transcription termination in kinetoplastids. Additional studies regarding the PJW/PP1 complex formation will help dissect the novel RNA Pol II transcription cycle in these divergent eukaryotes.

## Experimental Procedures

### Protein structure modeling with AlphaFold2

The predicted models were generated using the AlphaFold2 algorithm (73) via the ColabFold platform (74). In the open source Google CoLabFold platform, sequences were pasted in the query sequence box and the complex prediction was run with the default settings. The AlphaFold model was represented by five top-scored conformations along with estimates of prediction reliability (pLDDT), as described elsewhere (73). The protein models were analyzed and displayed with UCSF ChimeraX version: 1.5 (75).

### Parasite culture

Promastigote form *L. tarentolae* were grown in SDM79 medium at 27 L. Transfections were performed as previously described (53). After transfection, cells were plated into 96-well plates to obtain clonal cell lines by limiting dilution.

Where appropriate, the following drug concentrations were used: 50 g/ml G418 and 10 g/ml Puromycin. Bloodstream form *T. brucei* expressing T7 RNA polymerase and the Tet repressor (“single marker cells”) (76) were cultured in HMI- 9 medium at 37 ℃. Transfections were performed using the Amaxa electroporation system (Human T Cell Nucleofactor Kit, program X-001) and clonal cell lines obtained as described (53). Where appropriate, the following drug concentrations were used: 2ug/ml G418, 2.5 ug/ml Hygromycin, 2.5 ug/ml Phleomycin, 5 ug/ml Blasticidin, 0.2 ug/ml Puromycin, and 2 ug/ml Tetracycline.

### DNA constructs and cell line generation

Endogenous HA-tagging in *L. tarentolae*. A background *L. tarentolae* cell line was established to express Cas9 and T7 polymerase following transfection with PacI-digested pTB007 plasmid (59) as previously described (55). To tag the endogenous PP1-8e, PP1-7d, PNUTS, JBP3 or Wdr82 locus with 6xHA tag, the Cas9/T7-expressing cell line was transfected with gRNAs and donor fragments, as previously described (55). For overexpressing C-terminal tagged proteins in *L. tarentolae*, the open reading frame of LtPNUTS or LtPP1 was PCR amplified without a stop codon and inserted into the pSNSAP1 vector at the BamH1 and Xba1 sites as previously described (55). The obtained constructs were referred to as PNUTS-Pd or PP1-Pd. The desired PP1 or PNUTS mutants were generated by oligonucleotide-mediated site-directed mutagenesis (QuikChange II XL Site-Directed Mutagenesis Kit, Agilent Technologies) following the manufacturer’s instructions. All final constructs were sequenced prior to electroporation. PNUTS-Pd plasmid was transfected into the PP1-8e-HA cell line and WT *L. tarentolae* and the PP1-Pd plasmid transfected into the PNUTS-HA and WT cell line.

Endogenous tagging in *T. brucei*. For tagging the 3’ end of the TbPNUTS, Wdr82, and JBP3 with 3xHA or Myc tag in *T. brucei* cells, a PCR-based approach was used with the pMOTag4H or pMOTag3M vectors as described(53). For tagging PNUTS with the PTP tag in *T. brucei*, the 3’ end of TbPNUTS was cloned in the ApaI and Not1 sites of the Pc-PTP-Neo vector (77) where the Neomycin resistance drug marker was replaced with a blasticidin resistance drug marker. The vector was then linearized by restriction enzyme digestion within the 3’ end of the TbPNUTS gene, and used in transfection. For tetracycline regulated expression of PNUTS in *T. brucei*, the ORF with a C-terminal PTP tag was amplified by PCR and cloned into the HindIII and BamH1 sites of the pLew100V5 plasmid. The final construct (PNUTS-PTP-pLew100), was linearized with NotI prior to transfection. To induce PNUTS expression, tetracycline was added at 2 ug/ml. All final constructs were sequenced prior to electroporation. Primers sequences used are available upon request.

### RNAi analysis

Conditional silencing of PNUTS, JBP3 and Wdr82 in *T. brucei* BF SMC was performed as previously described (53).

### Co-immunoprecipitation

5 × 10^8^ of *L. tarentolae* cells were lysed in lysis buffer and Pd-tagged protein was affinity purified using 50 ul IgG Sepharose beads as previously described (37). After incubation with cell extract for 4 hrs at 4 L, the IgG beads were washed 3 times in 10 ml PA-150 buffer. After the final wash, the beads were boiled for 5 min in 1x SDS-PAGE sample buffer. Samples were run on 10% sodium dodecyl sulfate polyacrylamide gel and transferred to nitrocellulose membrane for western blotting with anti-protein A and anti-HA antibodies. 1.2 x 10^8^ of *T. brucei* cells expressing PTP-tagged protein was used for co-immunoprecipitation as described above. Samples were run on 10% sodium dodecyl sulfate polyacrylamide gel and transferred to nitrocellulose membrane for western blotting with anti-protein A, anti-HA and Anti-Myc antibodies.

### Western blotting

Proteins from 1.4 x 10^7^ cell equivalents of *L. tarentolae*, or 3 x 10^6^ of *T. brucei* cells, were separated by sodium dodecyl sulphate polyacrylamide gel electrophoresis (SDS page 10% gel), transferred to nitrocellulose and probed with anti-protein A (1:5000), anti-MYC (Santa Cruz, 9E10, 1:1000), anti-HA antibodies (Sigma, 3F10, 1:3000) or anti-Elongation Factor 1A (Sigma, 05-235, 1:20000). Bound antibodies were detected by an Alexa Fluor 680 labelled secondary goat anti-rat antibody, an Alexa Fluor 680 labelled secondary goat anti-mouse (LiCor), or an Alexa Fluor 800 labelled secondary goat ant-rabbit, and analyzed with Image Studio software (LiCor).

### Strand-specific RT-qPCR

Total RNA was extracted using the Tripure Isolation Reagent (Roche). To synthesize cDNA, 1 ug of Turbo™ DNase- treated total RNA (ThermoFisher) was reverse-transcribed with strand-specific oligonucleotides using Superscript™ III kit (ThermoFisher), following the manufacturer’s instructions. Quantification of the resulting cDNA was conducted using an iCycler with an iQ5 multicolor real-time PCR detection system (Bio-Rad). Triplicate cDNA samples were assessed and normalized against tubulin cDNA. For the qPCR reaction, a 15 ul mixture containing 5 ul of cDNA, 4.5 pmol each of sense and antisense primers, and 7.5 ul of 2× iQ SYBR green super mix (Bio-Rad Laboratories) was used.

Standard curves were generated for each gene using 5-fold dilutions of a known quantity (100 ng/l) of WT gDNA. The quantities were determined using the iQ5 optical detection system software.

## Data availability

All data described in this study are presented in the article and accompanying supporting information.

## Supporting information

This article contains supporting information.

