## supplemental data for "Leishmania PNUTS discriminates between PP1 catalytic subunits through a RVxF-ΦΦ-F motif and polymorphisms in the PP1 C-tail and catalytic domain"

**Figure S1**

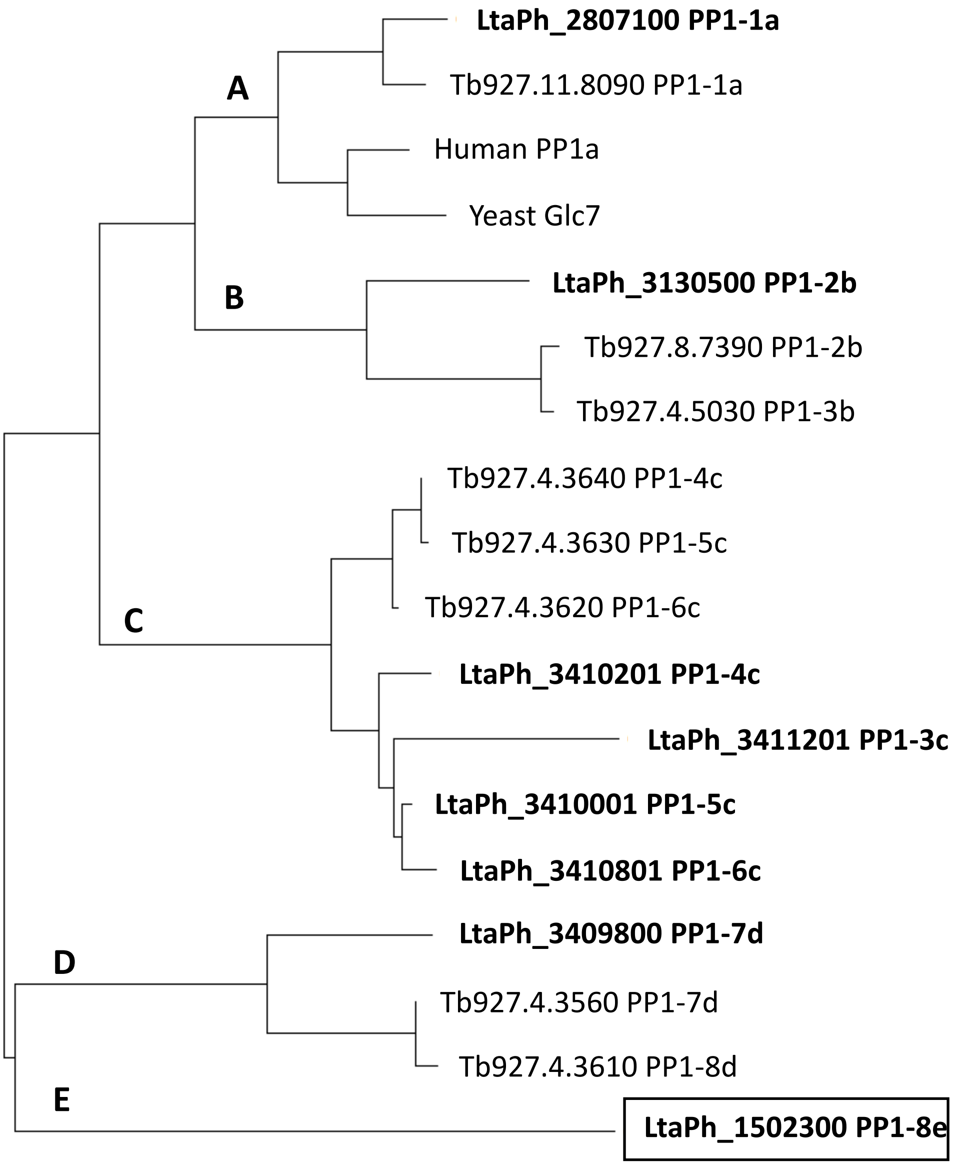


**Figure S1.** Phylogenetic analysis of PP1 isotypes in *L. tarentolae* (bold), *T. brucei*, human (PP1α) and yeast (Glc7). The tree was obtained with Maximum Likelihood method and JTT matrix-based model, using MEGA11 software. The five clades of PP1 (A-E) are shown. Genes for proteins used in the alignment for *L.* tarentolae and *T. brucei* are indicated. Glc7, *Saccharomyces cerevisiae* (SGD:S000000935).

### Figure S2

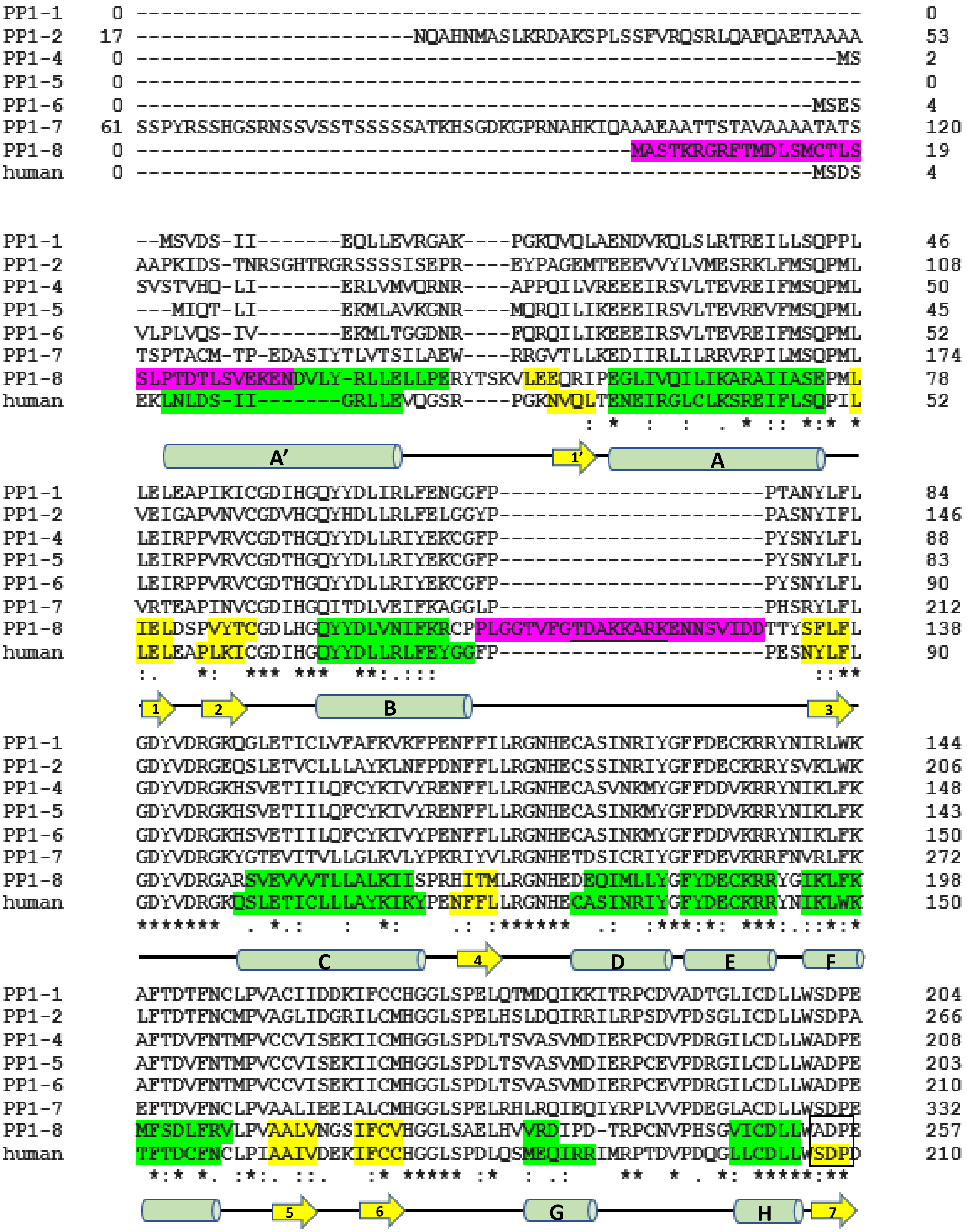
PP1-1 0 ------------------------------------------------------------------------- 0
PP1-2 17 -----------------------NQAHNMASLKRDAKSPLSSFVRQSRLQAFQAETAAAA 53
PP1-4 0 ----------------------------------------------------------MS 2
PP1-5 0 ------------------------------------------------------------ 0
PP1-6 0 --------------------------------------------------------MSES 4
PP1-7 61 SSPYRSSHGSRNSSVSSTSSSSSATKHSGDKGPRNAHKIQAAAEAATTSTAVAAAATATS 120
PP1-8 0 -----------------------------------------MASTKRGRFTMDLSMCTLS 19
human 0 --------------------------------------------------------MSDS 4
PP1-1 --MSVDS-II-------EQLLEVRGAK----PGKQVQLAENDVKQLSLRTREILLSQPPL 46
PP1-2 AAPKIDS-TNRSGHTRGRSSSSISEPR----EYPAGEMTEEEVVYLVMESRKLFMSQPML 108
PP1-4 SVSTVHQ-LI-------ERLVMVQRNR----APPQILVREEEIRSVLTEVREIFMSQPML 50
PP1-5 ---MIQT-LI-------EKMLAVKGNR----MQRQILIKEEEIRSVLTEVREVFMSQPML 45
PP1-6 VLPLVQS-IV-------EKMLTGGDNR----FQRQILIKEEEIRSVLTEVREIFMSQPML 52
PP1-7 TSPTACM-TP-EDASIYTLVTSILAEW----RRGVTLLKEDIIRLILRRVRPILMSQPML 174
PP1-8 SLPTDTLSVEKENDVLY-RLLELLPERYTSKVLEEQRIPEGLIVQILIKARAIIASEPML 78
human EKLNLDS-II-------GRLLEVQGSR----PGKNVQLTENEIRGLCLKSREIFLSQPIL 52
: * : : . * :: *:* *
A’
1’
A
PP1-1 LELEAPIKICGDIHGQYYDLIRLFENGGFP----------------------PTANYLFL 84
PP1-2 VEIGAPVNVCGDVHGQYHDLLRLFELGGYP----------------------PASNYIFL 146
PP1-4 LEIRPPVRVCGDTHGQYYDLLRIYEKCGFP----------------------PYSNYLFL 88
PP1-5 LEIRPPVRVCGDTHGQYYDLLRIYEKCGFP----------------------PYSNYLFL 83
PP1-6 LEIRPPVRVCGDTHGQYYDLLRIYEKCGFP----------------------PYSNYLFL 90
PP1-7 VRTEAPINVCGDIHGQITDLVEIFKAGGLP----------------------PHSRYLFL 212
PP1-8 IELDSPVYTCGDLHGQYYDLVNIFKRCPPLGGTVFGTDAKKARKENNSVIDDTTYSFLFL 138
human LELEAPLKICGDIHGQYYDLLRLFEYGGFP----------------------PESNYLFL 90
:. *: *** *** **:.::: ::**
1
2
B
3
PP1-1 GDYVDRGKQGLETICLVFAFKVKFPENFFILRGNHECASINRIYGFFDECKRRYNIRLWK 144
PP1-2 GDYVDRGEQSLETVCLLLAYKLNFPDNFFLLRGNHECSSINRIYGFFDECKRRYSVKLWK 206
PP1-4 GDYVDRGKHSVETIILQFCYKIVYRENFFLLRGNHECASVNKMYGFFDDVKRRYNIKLFK 148
PP1-5 GDYVDRGKHSVETIILQFCYKIVYRENFFLLRGNHECASINKMYGFFDDVKRRYNIKLFK 143
PP1-6 GDYVDRGKHSVETIILQFCYKIVYPENFFLLRGNHECASINKMYGFFDDVKRRYNIKLFK 150
PP1-7 GDYVDRGKYGTEVITVLLGLKVLYPKRIYVLRGNHETDSICRIYGFFDEVKRRFNVRLFK 272
PP1-8 GDYVDRGARSVEVVVTLLALKIISPRHITMLRGNHEDEQIMLLYGFYDECKRRYGIKLFK 198
human GDYVDRGKQSLETICLLLAYKIKYPENFFLLRGNHECASINRIYGFYDECKRRYNIKLWK 150
******* . *.: : *: .: :****** .: :***:*: ***:.::*:*
C
4
D
E
F
PP1-1 AFTDTFNCLPVACIIDDKIFCCHGGLSPELQTMDQIKKITRPCDVADTGLICDLLWSDPE 204
PP1-2 LFTDTFNCMPVAGLIDGRILCMHGGLSPELHSLDQIRRILRPSDVPDSGLICDLLWSDPA 266
PP1-4 AFTDVFNTMPVCCVISEKIICMHGGLSPDLTSVASVMDIERPCDVPDRGILCDLLWADPE 208
PP1-5 AFTDVFNTMPVCCVISEKIICMHGGLSPDLTSVASVMDIERPCEVPDRGILCDLLWADPE 203
PP1-6 AFTDVFNTMPVCCVISEKIICMHGGLSPDLTSVASVMDIERPCEVPDRGILCDLLWADPE 210
PP1-7 EFTDVFNCLPVAALIEEIALCMHGGLSPELRHLRQIEQIYRPLVVPDEGLACDLLWSDPE 332
PP1-8 MFSDLFRVLPVAALVNGSIFCVHGGLSAELHVVRDIPD-TRPCNVPHSGVICDLLWADPE 257
human TFTDCFNCLPIAAIVDEKIFCCHGGLSPDLQSMEQIRRIMRPTDVPDQGLLCDLLWSDPD 210
*:* *. :*:. ::. :* ***** :* : .: ** * . *: *****:**
5
6
G
H
7

```
PP1-1     E-GL--SGWGENDRGVSFTFGQDIVEKFLNKHQLELICRAHQVVEDGYQFFA---KRKLI      258
PP1-2     EDPI--TGFGENDRGVSWTFGENVVENITEALDLDLICRAHQVAEEGYKFFA---KRKLL      321
PP1-4     D-DV--QGFLESDRGVSYLFGEDIVNDFLDMVDMDLVVRAHQVMERGYGFFA---SRQLV      262
PP1-5     D-EV--RGFLESDRGVSYLFGEDIVNDFLDMVDMDLIVRAHQVVQRGYEFFA---SRQLV      257
PP1-6     D-EV--RGFLESDRGVSYLFGEDIVNDFLDMVDMDLIVRAHQVVQRGYEFFA---SRQLV      264
PP1-7     E-GT--SGWQPSERGVSFTFGEDVVKRMCDSLGIDIVLRAHQVVDEGYSFFA---GRRLV      386
PP1-8     TDLPAGVDWAPSSRRISSVFSERALERFLVENDIDLVCRAHQVVEEGFQFFPDHKKRHLL      317
human     K-DV—QGWGENDRGVSFTFGAEVVAKFLHKHDLDLICRAHQVVEDGYEFFA---KRQLV      264
               .:   ..* :*   *.    :   :     :::: ***** : *: **     *:*:
```

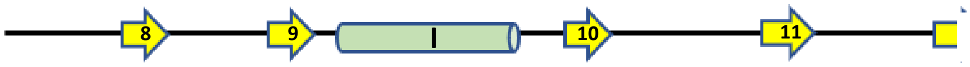


```
PP1-1     TIFSAPNYCNEFDNSGAVMTVDNELMCSFQILKPSVKKPKFYS----------------     301
PP1-2     TVFSAPNYCGEFNNYGAFLCVDENLMCSVKQLVPLFEVNDFECE---------------     365
PP1-4     TVFSAPNYCGEFDNDAAVMTIDDKLQCSFLIIPAFR-----------------------     298
PP1-5     TVFSAPNYCGEFDNDAAVMTIDDKLQCSFLIIPARK-----------------------     293
PP1-6     TVFSAPNYCGEFDNDAAVMTIDDKLQCSFLIVPAPH-----------------------     300
PP1-7     TIFSASNYCGEFTNSGAMMLMDENCMCSFQIFKPEY-----------------------     422
PP1-8     TIFSATNYCNEFGNRGGILRVDEKGVCSIITLEPPDFVQQRDIMLLRDPLPNPLSHD---     374
human     TLFSAPNYCGEFDNAGAMMSVDETLMCSFQILKPADKNKGKYGQFSGLNPGGRPITPPRN     324
          *:*** ***.** *  ...: :*:    **.   .
```

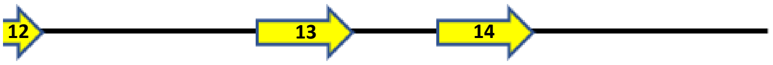


```
PP1-1     ------          301
PP1-2     ------          365
PP1-4     ------          298
PP1-5     ------          293
PP1-6     ------          300
PP1-7     ------          422
PP1-8     ------          374
human     SAKAKK          330
```

**Figure S2.** Sequence alignment of *L. tarentolae* PP1 isoforms and human PP1α. Identical (*), conservative (:) and similar (.) residues between hPP1 and LtPP1-8 are indicated. Cylinders and arrows representing α-helices and β-strands of the hPP1α catalytic domain, respectively, are drawn under the sequence. The sequences corresponding to α-helices and β-strands are highlighted in the hPP1α sequence in green and yellow, respectively. The predicted secondary structure elements of Lt PP1-8e based on AlphaFold are also highlighted in green and yellow. The box at β-sheet 7 indicates that alanine substitution in the four amino acid element in LtPP1-8e leads to the inability of AlphaFold to predict a conserved β-sheet 7. But all the other α-helices and β-sheets within the catalytic domain are predicted. The distinct N- and C-termini and insertions within the catalytic domain of LtPP1-8e are highlighted in purple. Predicted α-helices within the 26-aa insert and C-terminus of LtPP1-8e are indicated by the underlined residues.

### Figure S3

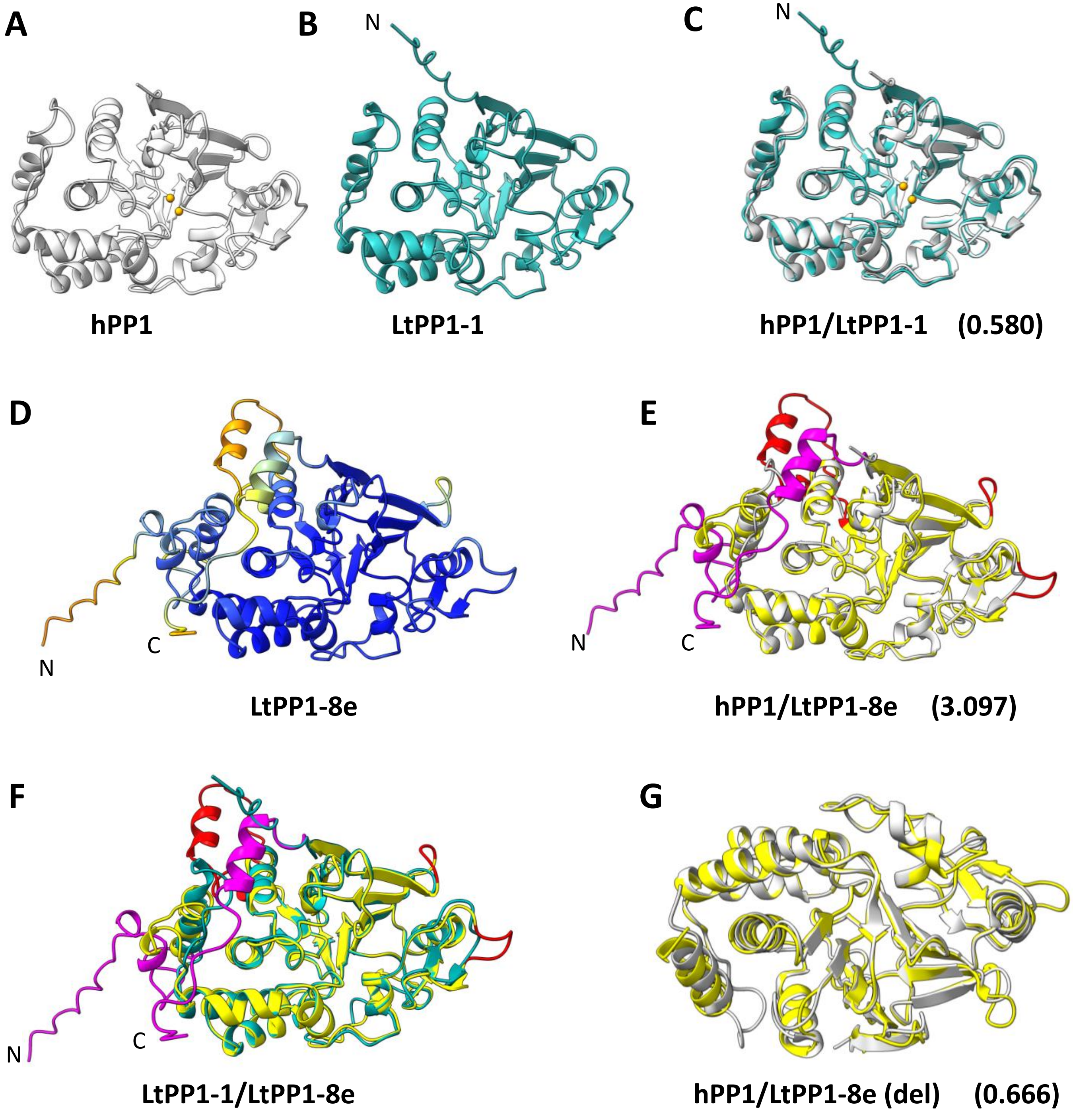


**Figure S3.** Structural homology of *L. tarentolae* PP1-8e and Human PP1α catalytic domains. A, Human PP1α (7-300) structure (3E7A) shown as a carton in grey. Bound metal ions are shown as orange spheres. B, Predicted LtPP1-1a structure illustrated as a carton in blue. C, Structural overlay of hPP1 structure and predicted LtPP1-1a. The root-mean-square-deviation (RMSD), estimating the degree of structural similarity between the model and the crystal data, is indicated. D, The predicted Lt PP1-8e structure colored by pLDDT values, with blue representing high model confidence (pLDDT > 90) and orange representing low model confidence (70 > pLDDT > 50). E, Structural overlay of hPP1 (grey) and LtPP1-8e. LtPP1-8e is in yellow with unique sequences within the catalytic domain and the extremities are highlighted in red and magenta, respectively. F, Structural overlay of LtPP1-1a and LtPP1-8e models with the PP1-8e unique sequences colored as in E. G, Structural overlay of hPP1 structure (grey) and LtPP1-8e model with the unique sequences within the catalytic motif and extremities deleted (del).

### Figure S4

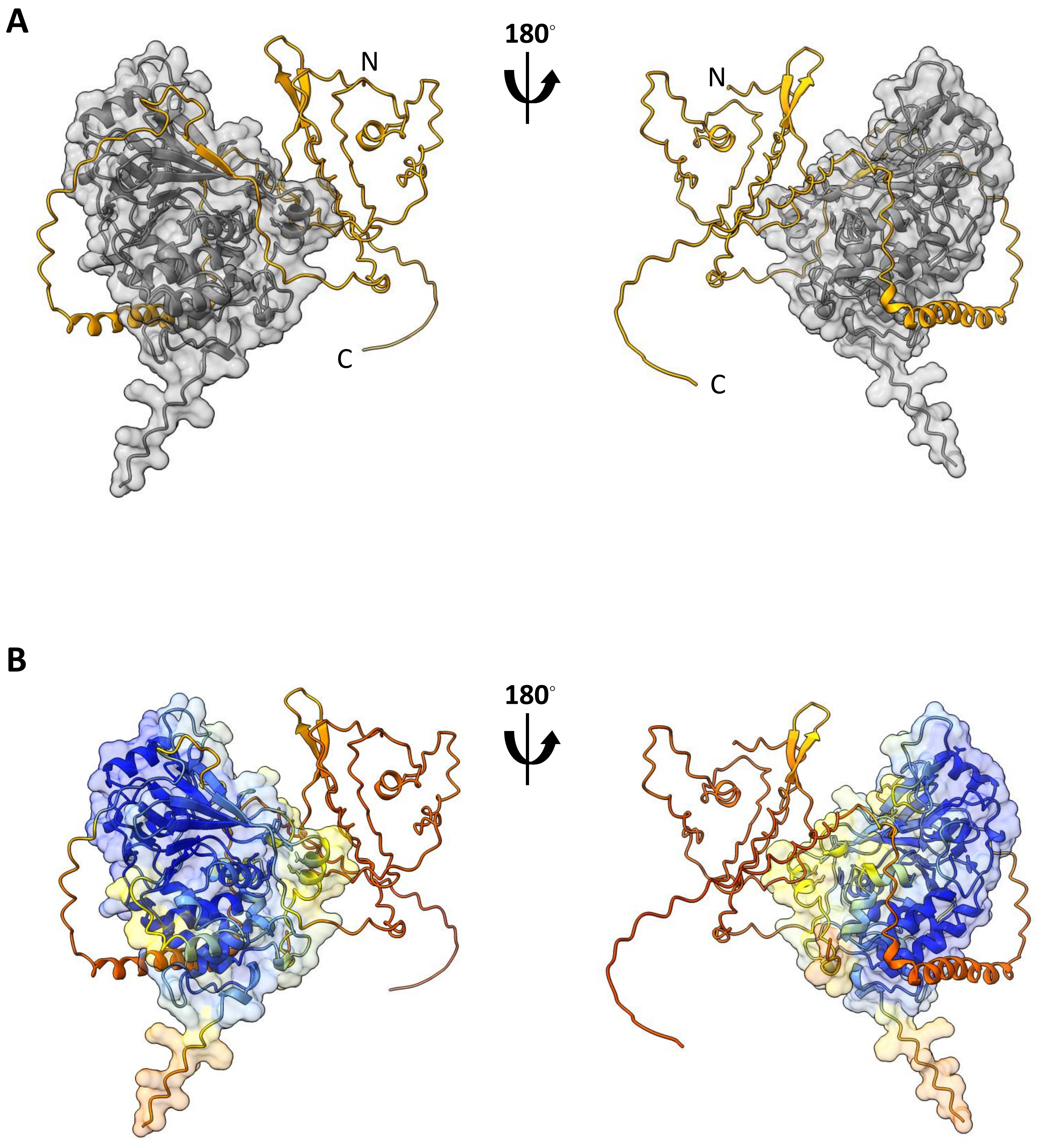


**Figure S4.** AlphaFold model of the Lt PP1-8e:PNUTS complex.  A, The predicted LtPNUTS (orange cartoon) and LtPP1-8e (gray surface) complex. B, The predicted LtPNUTS:LtPP1-8e complex colored by pLDDT values, with blue representing high model confidence (pLDDT > 90) and orange representing low model confidence (70 > pLDDT > 50).

### Figure S5

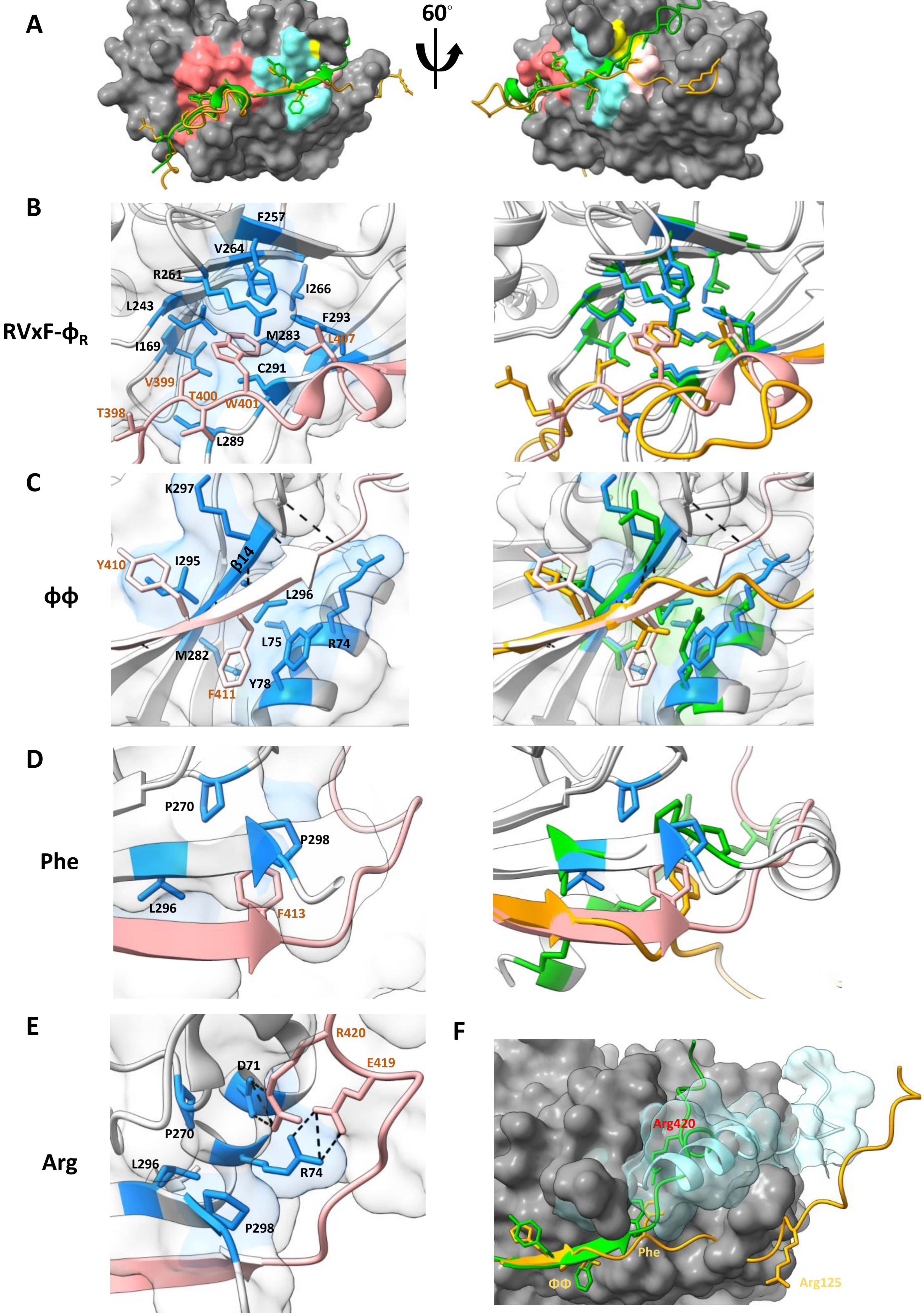

A
60°
B
RVxF-ϕR
F257
V264
R261
I266
L243
F293
M283
L407
I169
C291
V399
T400
W401
T398
L289
C
ϕϕ
K297
Y410
I295
β14
L296
L75
R74
M282
Y78
F411
D
Phe
P270
P298
L296
F413
E
Arg
R420
E419
D71
P270
R74
L296
P298
F
Arg420
Phe
ΦΦ
Arg125

### Figure S5 (continued)

**G** **H**

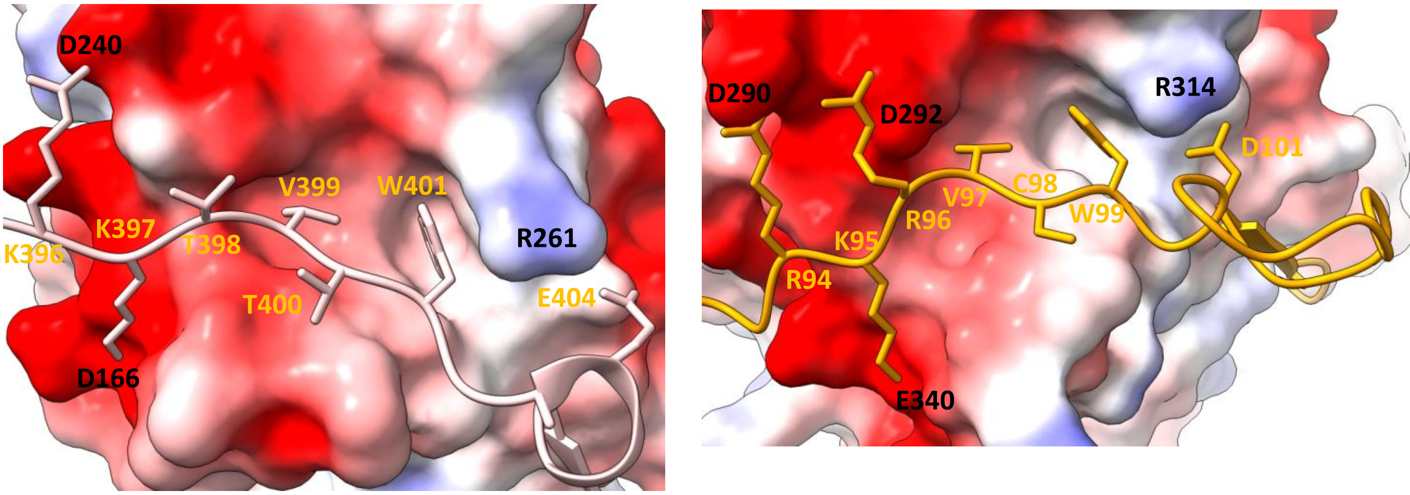


**hPP1-PNUTS** **LtPP1-PNUTS**

```
                       RVxF               ΦR
LtPNUTS   91APSRKRVCWADEGHTDVSRGLVKHVTNFYMPNT.SR125
 hPNUTS  393RKRKKTVTWPEEGK.......LREYF.YFELDET.ER420
```

**Figure S5.** LtPNUTS and hPNUTS share PP1-binding motifs. A, The predicted LtPNUTS:PP1-8e structure is superimposed to the human PP1:PNUTS structure (4mp0). Human PP1 is shown as grey surface. LtPNUTS (orange) and hPNUTS (green) are represented as ribbons with key interacting residues shown as sticks. The RVxF binding pocket (red), ΦΦ binding pocket (cyan), Phe binding pocket (yellow) and Arg binding pocket (pink) are shaded on the hPP1 surface and correspond to the zoomed-in pockets shown in B, C, D and E. Close-up of the RVxF binding pocket (B), ΦΦ binding pocket (C), and Phe (D) and Arg binding pockets (E) on the human PP1:PNUTS holoenzyme. B-D; Left, key interacting residues in hPP1 (blue) and hPNUTS (pink) are shown as sticks and labelled. Right, overlay of the Lt and human PP1-PNUTS structures, with key interactive residues between LtPP1-8e (green) and LtPNUTS (orange) shown as sticks. E, Arg binding pocket on the human PP1:PNUTS holoenzyme. Salt bridge interactions between PNUTS R420 and E419 with the corresponding residues of the Arg binding pocket in PP1 are indicated by the dashed line. F, structural overlay of the human PP1-PNUTS structure and Lt PP1-PNUTS model. hPNUTS (green) is bound to hPP1 (grey surface) with Arg420 shown as stick. Arg125 in LtPNUTS is not predicted to bind into the Arg binding pocket. The Arg binding pocket in the human structure is occupied by the C-terminus of PP1-8e (shown as blue ribbon and surface). G and H, electrostatic surface potential representation (positive, blue; negative, red) of the RVxF-Or pocket in the hPNUTS:PP1 complex (G) and LtPNUTS:PP1-8e complex (H). The sequence alignments for these regions of PNUTS are provided below.

### Figure S6

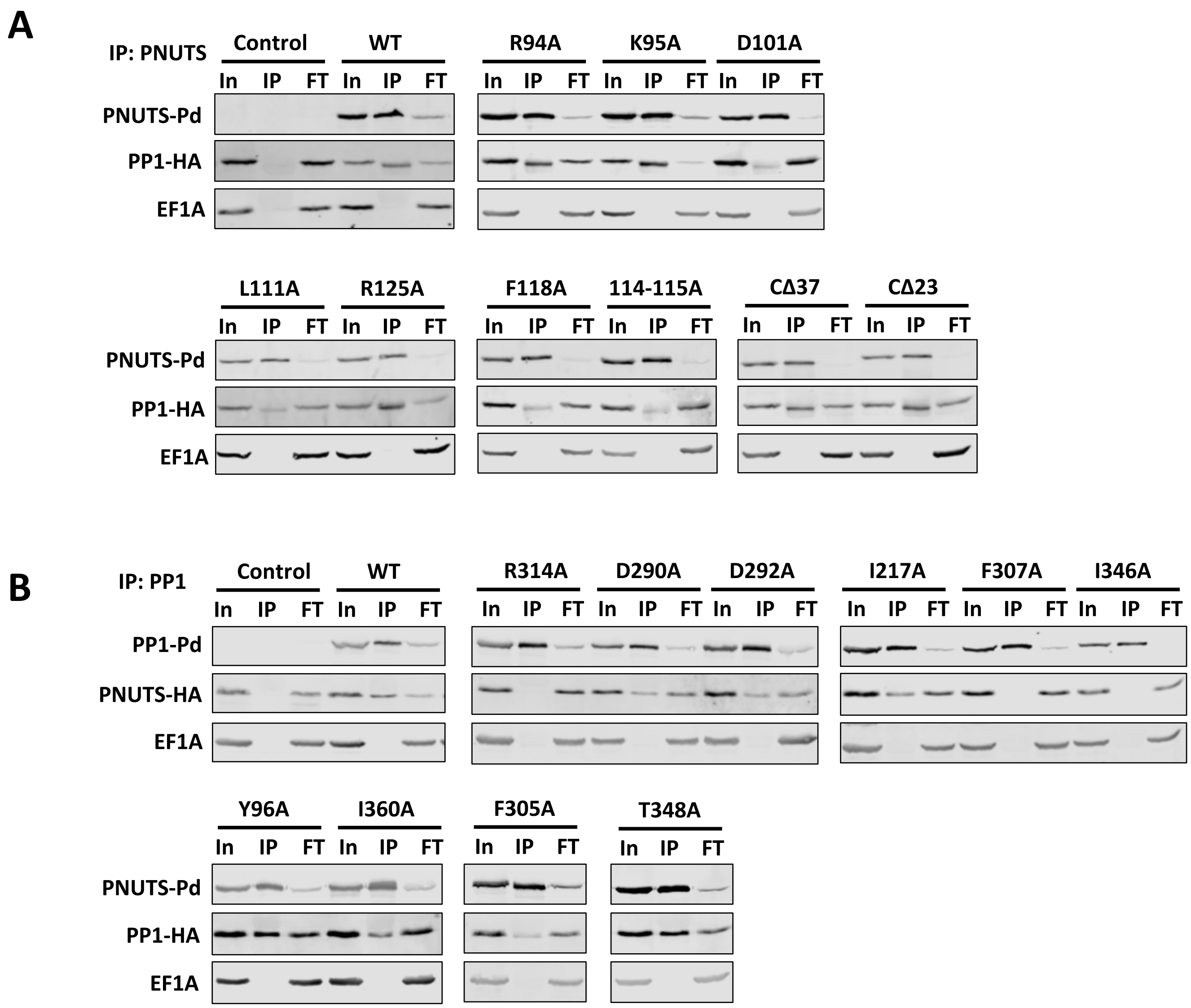


**Figure S6.** Co-immunoprecipitation analysis of LtPP1-PNUTS. Western blots showing the Co-IP of Pd-tagged wild type or indicated PNUTS (A) or PP1 (B) mutants with HA-tagged PP1 or PNUTS, respectively, as described in Figure 3. A representative gel image from three independent experiments for each Co-IP is shown.

### Figure S7

```
PP1-1     ------------------------------------------------------------    0
PP1-2     ---------------------------MSGLSKVEIIISKL-------------LL----    16
PP1-4     ------------------------------------------------------------    0
PP1-5     ------------------------------------------------------------    0
PP1-6     ------------------------------------------------------------    0
PP1-7     MRHGTTPQVGGRKRSRSHSSEKSPVPALHDEGAEDVVASSVHAKEKYHQQLSATVTSPAI    60
PP1-8     ------------------------------------------------------------    0
human     ------------------------------------------------------------    0


PP1-1     ------------------------------------------------------------    0
PP1-2     ----NQAHN-------------MASLKRDAKSPLSSFVRQSRLQAFQAET------AAAA    53
PP1-4     ----------------------------------------------------------MS    2
PP1-5     ------------------------------------------------------------    0
PP1-6     --------------------------------------------------------MSES    4
PP1-7     SSPYRSSHGSRNSSVSSTSSSSSATKHSGDKGPRNAHKIQAAAEAATTSTAVAAAATATS    120
PP1-8     ----------------------MASTKRGR---------------FTMDL----SMCTLS    19
human     --------------------------------------------------------MSDS    4


PP1-1     --MSVDS-II-------EQLLEVRGAK----PGKQVQLAENDVKQLSLRTREILLSQPPL    46
PP1-2     AAPKIDS-TNRSGHTRGRSSSSISEPR----EYPAGEMTEEEVVYLVMESRKLFMSQPML    108
PP1-4     SVSTVHQ-LI-------ERLVMVQRNR----APPQILVREEEIRSVLTEVREIFMSQPML    50
PP1-5     ---MIQT-LI-------EKMLAVKGNR----MQRQILIKEEEIRSVLTEVREVFMSQPML    45
PP1-6     VLPLVQS-IV-------EKMLTGGDNR----FQRQILIKEEEIRSVLTEVREIFMSQPML    52
PP1-7     TSPTACM-TP-EDASIYTLVTSILAEW----RRGVTLLKEDIIRLILRRVRPILMSQPML    174
PP1-8     SLPTDTLSVEKENDVLY-RLLELLPERYTSKVLEEQRIPEGLIVQILIKARAIIASEPML    78
human     EKLNLDS-II-------GRLLEVQGSR----PGKNVQLTENEIRGLCLKSREIFLSQPIL    52
                                                 : *  :  :  . * :: *:* *

PP1-1     LELEAPIKICGDIHGQYYDLIRLFENGGFP----------------------PTANYLFL    84
PP1-2     VEIGAPVNVCGDVHGQYHDLLRLFELGGYP----------------------PASNYIFL    146
PP1-4     LEIRPPVRVCGDTHGQYYDLLRIYEKCGFP----------------------PYSNYLFL    88
PP1-5     LEIRPPVRVCGDTHGQYYDLLRIYEKCGFP----------------------PYSNYLFL    83
PP1-6     LEIRPPVRVCGDTHGQYYDLLRIYEKCGFP----------------------PYSNYLFL    90
PP1-7     VRTEAPINVCGDIHGQITDLVEIFKAGGLP----------------------PHSRYLFL    212
PP1-8     IELDSPVYTCGDLHGQYYDLVNIFKRCPPLGGTVFGTDAKKARKENNSVIDDTTYSFLFL    138
human     LELEAPLKICGDIHGQYYDLLRLFEYGGFP----------------------PESNYLFL    90
          :.   *:  *** ***  **:.:::                               ::**
                            ▲  ▲

PP1-1     GDYVDRGKQGLETICLVFAFKVKFPENFFILRGNHECASINRIYGFFDECKRRYNIRLWK    144
PP1-2     GDYVDRGEQSLETVCLLLAYKLNFPDNFFLLRGNHECSSINRIYGFFDECKRRYSVKLWK    206
PP1-4     GDYVDRGKHSVETIILQFCYKIVYRENFFLLRGNHECASVNKMYGFFDDVKRRYNIKLFK    148
PP1-5     GDYVDRGKHSVETIILQFCYKIVYRENFFLLRGNHECASINKMYGFFDDVKRRYNIKLFK    143
PP1-6     GDYVDRGKHSVETIILQFCYKIVYPENFFLLRGNHECASINKMYGFFDDVKRRYNIKLFK    150
PP1-7     GDYVDRGKYGTEVITVLLGLKVLYPKRIYVLRGNHETDSICRIYGFFDEVKRRFNVRLFK    272
PP1-8     GDYVDRGARSVEVVVTLLALKIISPRHITMLRGNHEDEQIMLLYGFYDECKRRYGIKLFK    198
human     GDYVDRGKQSLETICLLLAYKIKYPENFFLLRGNHECASINRIYGFYDECKRRYNIKLWK    150
          *******  . *.:   :  *:    .: :******  .:  :***:*: ***:.::*:*

PP1-1     AFTDTFNCLPVACIIDDKIFCCHGGLSPELQTMDQIKKITRPCDVADTGLICDLLWSDPE    204
PP1-2     LFTDTFNCMPVAGLIDGRILCMHGGLSPELHSLDQIRRILRPSDVPDSGLICDLLWSDPA    266
PP1-4     AFTDVFNTMPVCCVISEKIICMHGGLSPDLTSVASVMDIERPCDVPDRGILCDLLWADPE    208
PP1-5     AFTDVFNTMPVCCVISEKIICMHGGLSPDLTSVASVMDIERPCEVPDRGILCDLLWADPE    203
PP1-6     AFTDVFNTMPVCCVISEKIICMHGGLSPDLTSVASVMDIERPCEVPDRGILCDLLWADPE    210
PP1-7     EFTDVFNCLPVAALIEEIALCMHGGLSPELRHLRQIEQIYRPLVVPDEGLACDLLWSDPE    332
PP1-8     MFSDLFRVLPVAALVNGSIFCVHGGLSAELHVVRDIPD-TRPCNVPHSGVICDLLWADPE    257
human     TFTDCFNCLPIAAIVDEKIFCCHGGLSPDLQSMEQIRRIMRPTDVPDQGLLCDLLWSDPD    210
           *:* *. :*:. ::.   :* ***** :*  : .:    **  * . *: *****:**
```

#### Figure S7 continued

```
PP1-1      E-GL--SGWGENDRGVSFTFGQDIVEKFLNKHQLELICRAHQVVEDGYQFFA---KRKLI   258
PP1-2      EDPI--TGFGENDRGVSWTFGENVVENITEALDLDLICRAHQVAEEGYKFFA---KRKLL   321
PP1-4      D-DV--QGFLESDRGVSYLFGEDIVNDFLDMVDMDLVVRAHQVMERGYGFFA---SRQLV   262
PP1-5      D-EV--RGFLESDRGVSYLFGEDIVNDFLDMVDMDLIVRAHQVVQRGYEFFA---SRQLV   257
PP1-6      D-EV--RGFLESDRGVSYLFGEDIVNDFLDMVDMDLIVRAHQVVQRGYEFFA---SRQLV   264
PP1-7      E-GT--SGWQPSERGVSFTFGEDVVKRMCDSLGIDIVLRAHQVVDEGYSFFA---GRRLV   386
PP1-8      TDLPAGVDWAPSSRRISSVFSERALERFLVENDIDLVCRAHQVVEEGFQFFPDHKKRHLL   317
human      K-DV--QGWGENDRGVSFTFGAEVVAKFLHKHDLDLICRAHQVVEDGYEFFA---KRQLV   264
                  .:  ..* :*  *.   :  :     :::: ***** : *: **     *:*:

PP1-1      TIFSAPNYCNEFDNSGAVMTVDNELMCSFQILKPSVKKPKFYS-----------------   301
PP1-2      TVFSAPNYCGEFNNYGAFLCVDENLMCSVKQLVPLFEVNDFECE----------------   365
PP1-4      TVFSAPNYCGEFDNDAAVMTIDDKLQCSFLIIPAFR------------------------   298
PP1-5      TVFSAPNYCGEFDNDAAVMTIDDKLQCSFLIIPARK------------------------   293
PP1-6      TVFSAPNYCGEFDNDAAVMTIDDKLQCSFLIVPAPH------------------------   300
PP1-7      TIFSASNYCGEFTNSGAMMLMDENCMCSFQIFKPEY------------------------   422
PP1-8      TIFSATNYCNEFGNRGGILRVDEKGVCSIITLEPPDFVQQRDIMLL-----RDPLPNPLS   372
human      TLFSAPNYCGEFDNAGAMMSVDETLMCSFQILKPADKNKGKYGQFSGLNPGGRPITPPRN   324
           *:*** ***.** * ...: :*:   **.  .
                ▲                         ▲ ▲

PP1-1      ------        301
PP1-2      ------        365
PP1-4      ------        298
PP1-5      ------        293
PP1-6      ------        300
PP1-7      ------        422
PP1-8      HD----        374
human      SAKAKK        330
```

**Figure S7.** Sequence alignment of *L. tarentolae* PP1 isoforms and human PP1a. Identical (*), conservative (:) and similar (.) residues are indicated. Residues that form the extended RVxF binding pocket (yellow), ϕϕ binding pocket (purple), and Phe binding pocket (green) in the hPNUTS:hPP1 structure and predicted LtPNUTS:PP1-8e structure are highlighted. Residue that is a component of both the ϕϕ binding pocket and Phe binding pocket is highlighted in red. Residues that form the Arg binding pocket in hPP1 are indicated by a triangle underneath. Conserved residues among the remaining isoforms are also highlighted. Alignment was generated with ClustalW.

Figure S8

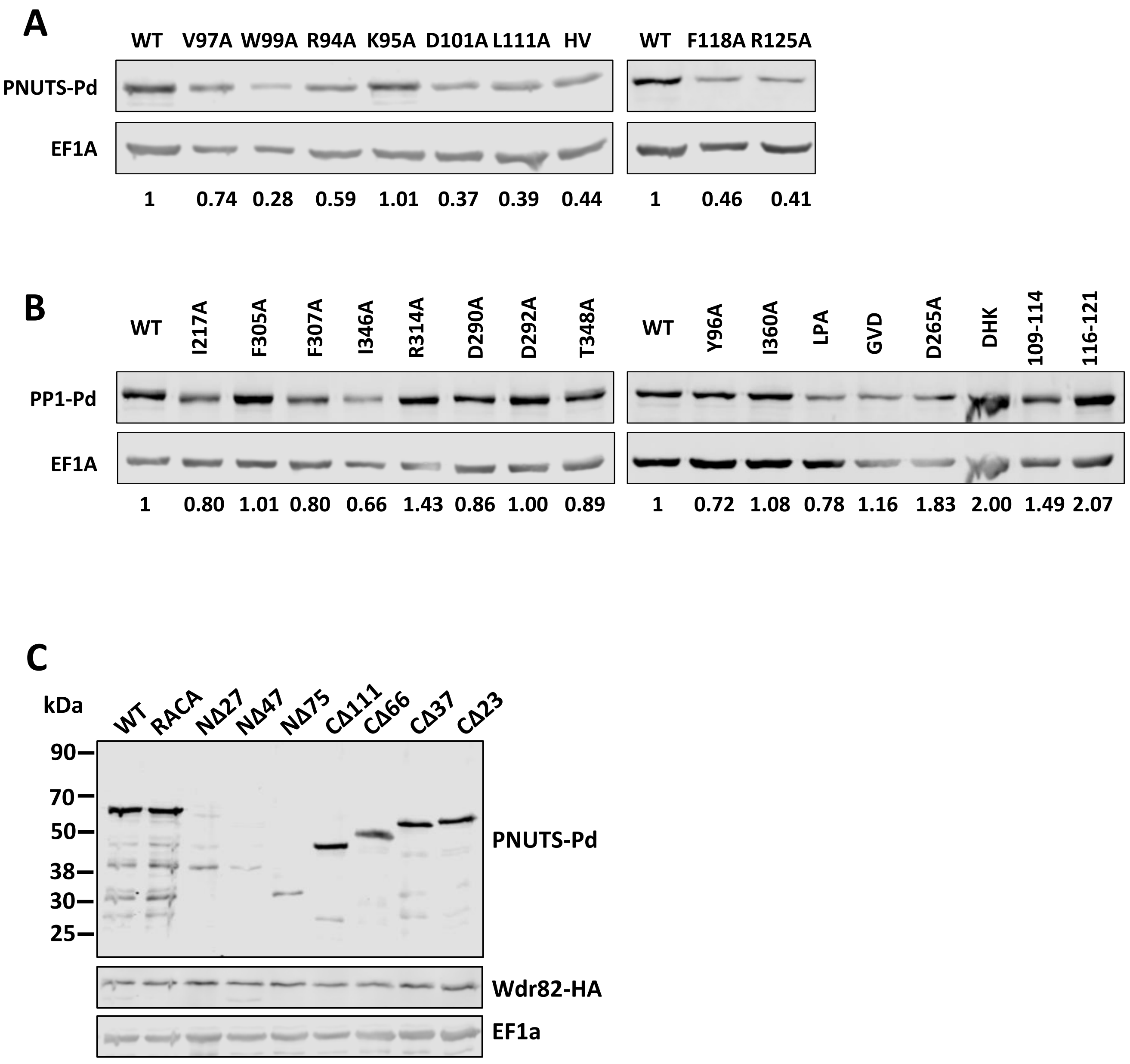


**Figure S8.** Expression levels of PNUTS and PP1. Lysates from cells that over-express the indicated PNUTS (A and C) or PP1 (B) mutant were analyzed by western blot with anti-protein A and anti-EF1A. EF1A serves as a loading control. Bands were quantified by densitometry and protein expression for the indicated mutant were normalized to EF1A. The normalized values are shown below the gel with wild type set to 1.

**Figure S9**

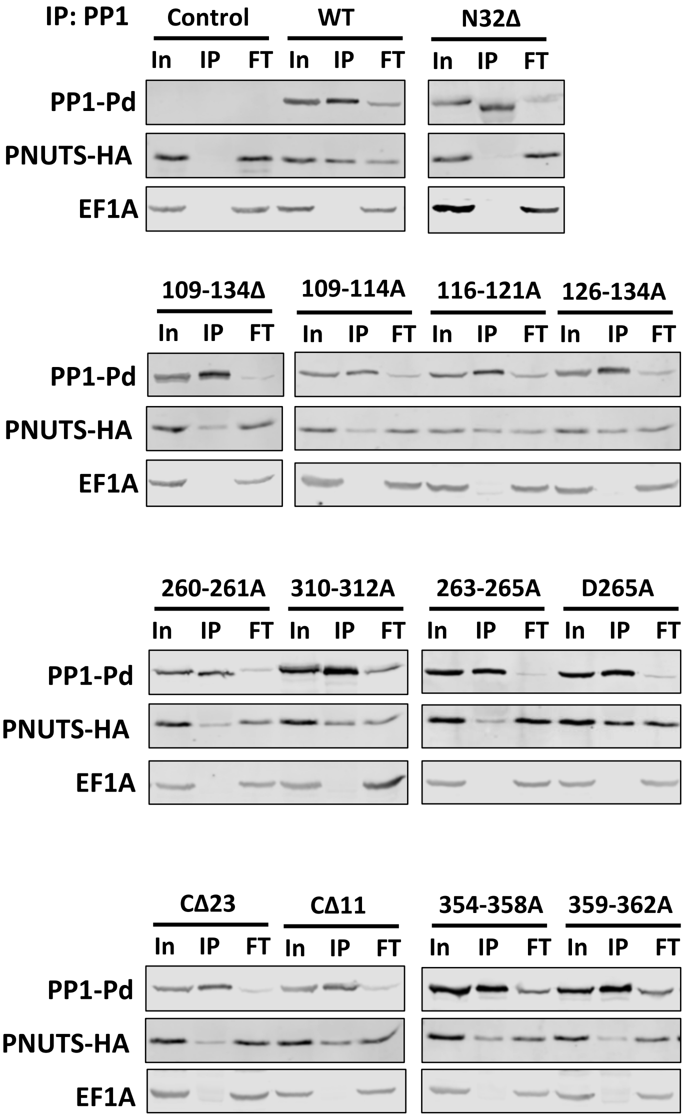


**Figure S9.** Coimmunoprecipitation analysis of LtPP1-PNUTS. Western blots showing the Co-IP of Pd-tagged PP1-8e (WT and indicated variants) with PNUTS-HA as described in Figure 4. A representative gel image from three independent experiments for each Co-IP is shown.

**Figure S10**

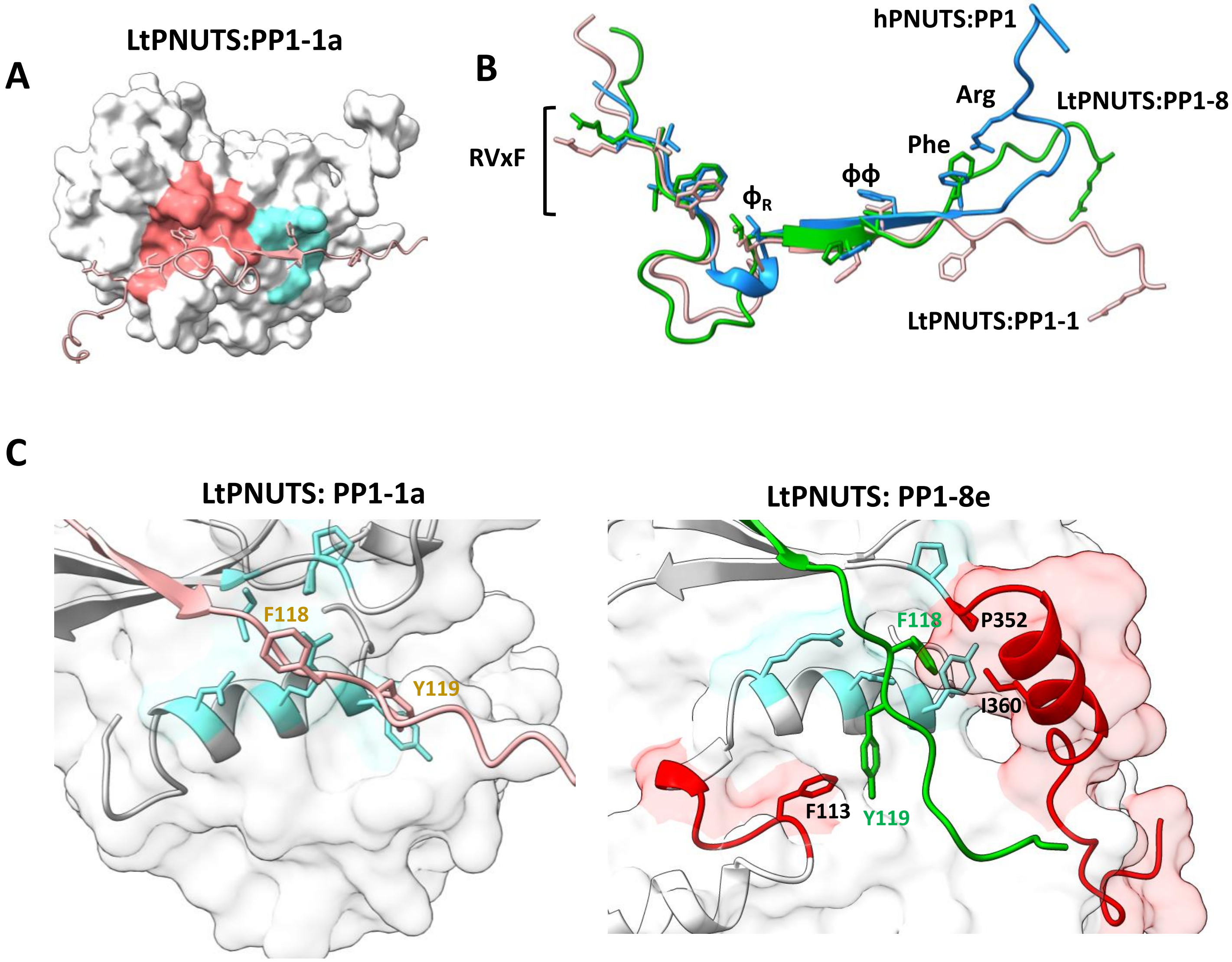


**Figure S10**. The predicted LtPNUTS-PP1-1a structure. A, The predicted holoenzyme structure of LtPNUTS (pink ribbon) and LtPP1-1a (white surface). The RVxF binding pocket (red) and ΦΦ binding pocket (cyan) are shaded on PP1 surface. B, Structural comparison of the PP1-binding domains of LtPNUTS in complex with PP1-8e (green) or PP1-1a (pink). Structure of hPNUTS (blue) bound to hPP1 is also shown. C, Close-up view of $F118_{PNUTS}$ (pink) in complex with LtPP1-1a (white surface, left) or (green) with LtPP1-8e (white surface, right). F118 and Y119 of PNUTS are shown as sticks and labelled. Conserved Phe-binding pocket residues (according the hPNUTS:PP1 structure) are shown in blue sticks. LtPNUTS:PP1-8e complex (right). The C-terminus of LtPP1-8e is shown in red carton, and key residues (P352 and I360) shown to be important for LtPNUTS binding are shown in sticks and labelled. Residues $^{109}$GGTVFG$^{114}$ within the PP1 catalytic motif and important for PNUTS binding are also shown in red, and residue $^{113}$F shown as sticks and labelled.

**Figure S11**

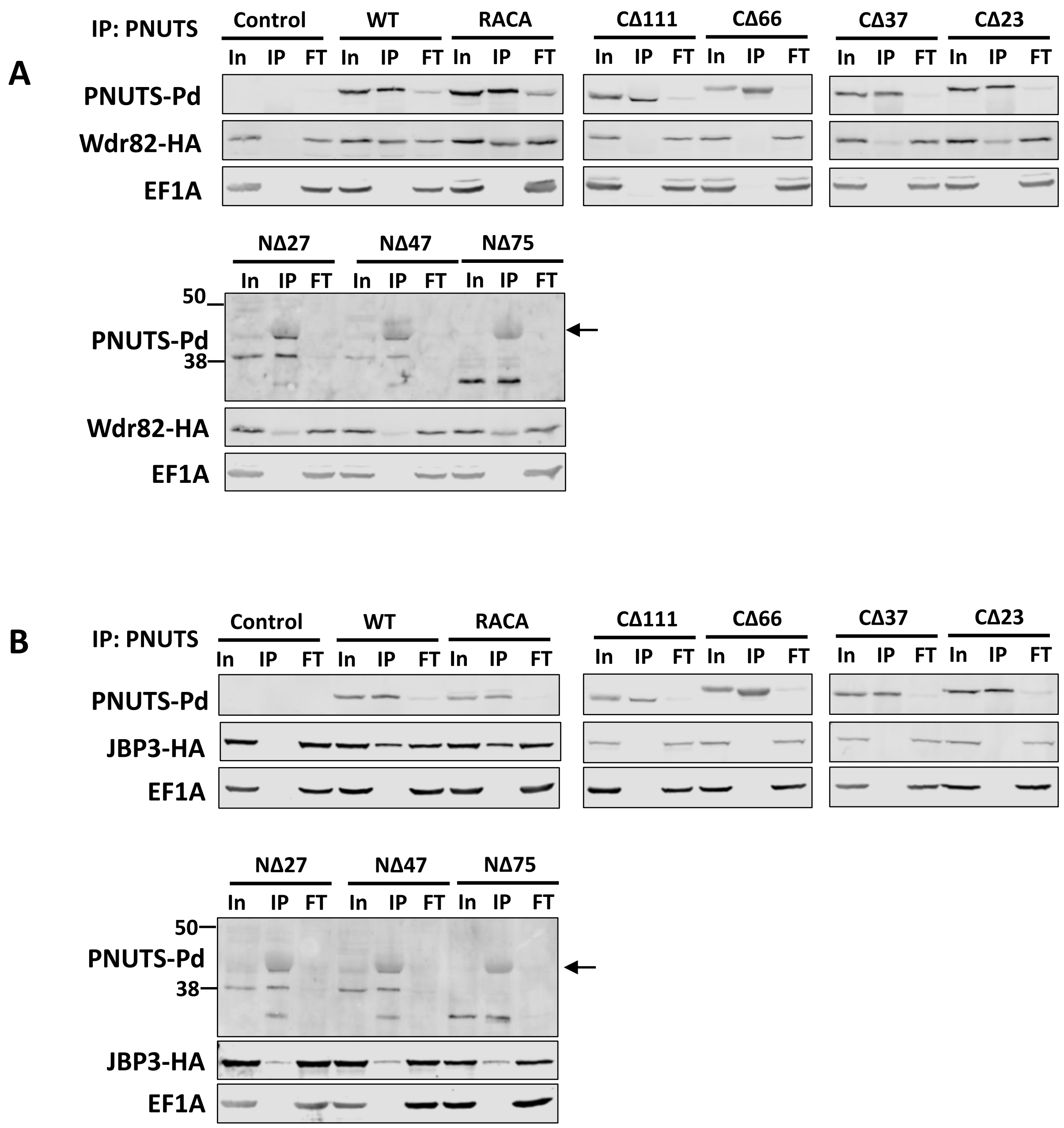


**Figure S11.** Coimmunoprecipitation analysis of LPNUTS and Wdr82 and JBP3. Pd-tagged PNUTS (WT or truncations) were expressed in Lt cells and tested for interaction with HA-tagged Wdr82 (A) or JBP3 (B) by Co-IP analysis as described in Figure 5. A representative Western blot image from three independent experiments for each Co-IP is shown. Arrow indicates IgG contamination in the IP of N-terminal PNUTS mutants.

**Figure S12**

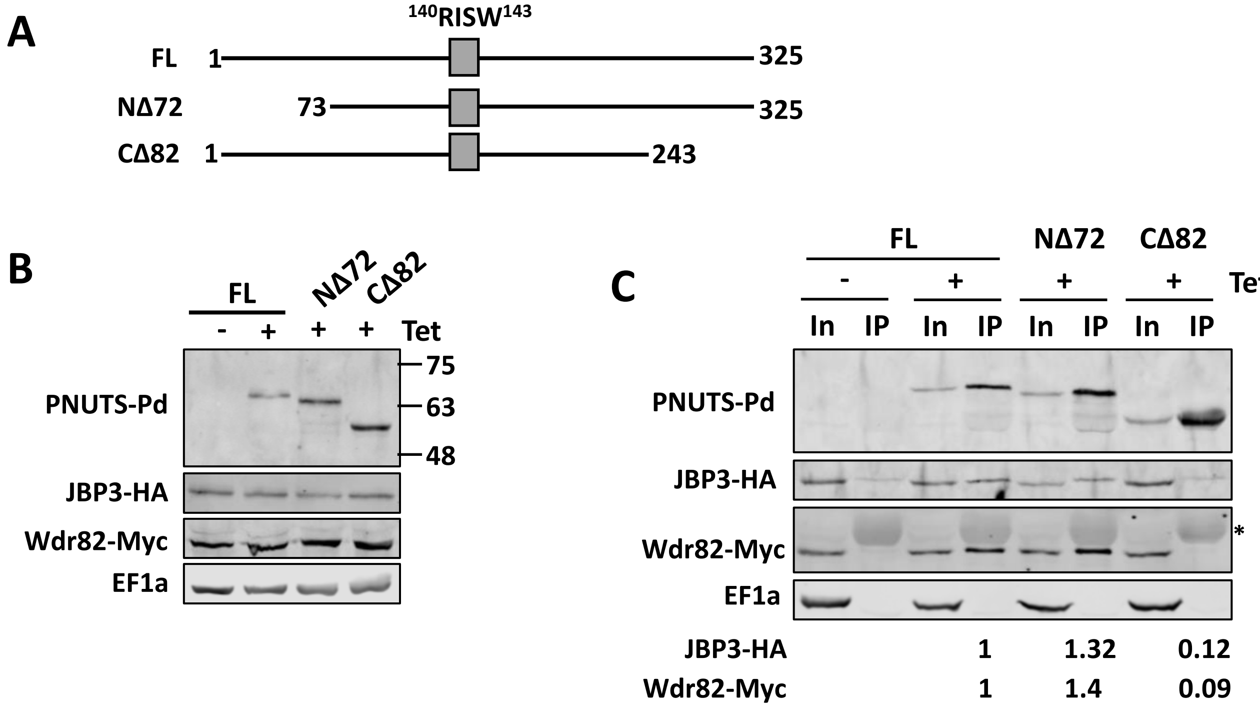


**Figure S12.** Co-immunoprecipitation analysis of TbPNUTS and Wdr82 and JBP3. A, Schematic representation of the TbPNUTS truncations. The putative RVXF motif is indicated by a grey box. B, JBP3 and Wdr82 were endogenously tagged with HA and Myc tags, respectively. The protein expression of the indicated TbPNUTS was induced by addition of tetracyclin (Tet) for 24 hrs and lysates analyzed by western blot with anti-protein A, anti-HA, anti-Myc, or anti-EF1a. EF1a serves as a loading control. C, Lysates of the indicated cell lines with or without tetracycline induction were purified by anti-protein A affinity resin and analyzed by western blot with anti-protein A, anti-HA, anti-Myc and anti-EF1a antibodies. FL, full-length (WT) TbPNUTS. Asterisk indicates the IgG cross-reactive signal in the IP fraction from anti-Myc. Protein A purification results in low background JBP3-HA signal in the absence of protein A-tagged PNUTS. %IP is quantified from two replicates and shown below for the corresponding cell lines.

Figure S13

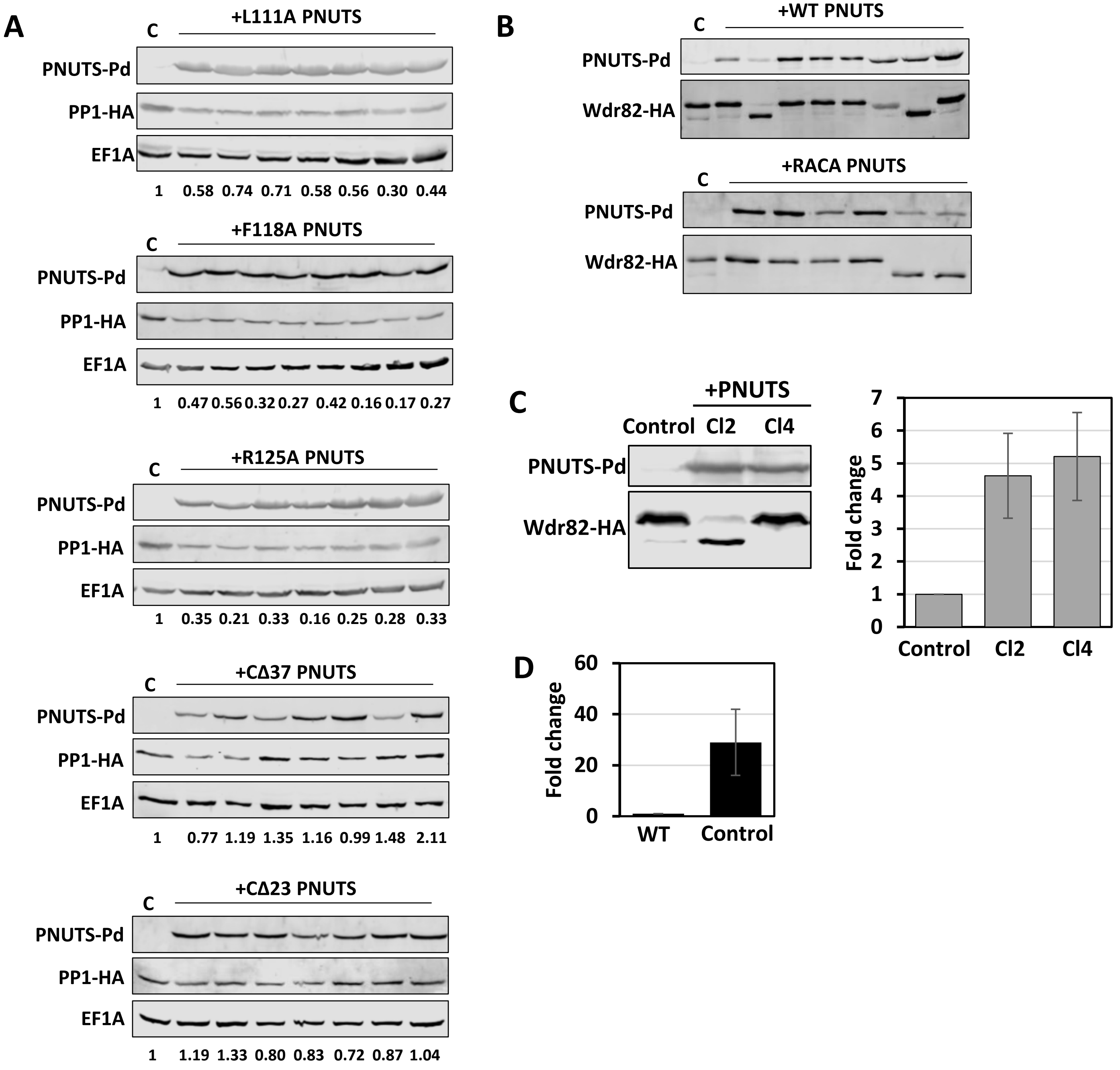


**Figure S13. Effects of PNUTS over-expression on PP1 and Wdr.** A, Effect of PNUTS over-expression on PP1-8e levels. LtPP1-8e was endogenously tagged with the HA tag and lysates from clonal cell lines following transfection of the indicated PNUTS construct were analyzed by western blot with anti-protein A and anti-HA. Anti-EF1A serves as a loading control. Bands were quantified by densitometry and PP1 levels were normalized to EF1A. The normalized values are shown below the gel with the untransfected PNUTS control cell set to 1. Control cell line lacking the PNUTS expression plasmid is indicated by the C. B, effect of PNUTS over-expression on Wdr82. Wdr82 was endogenously tagged with HA and cells transfected with the indicated PNUTS expression plasmid. Lysates from clonal cell lines were analyzed by western blot with anti-protein A and anti-HA. C, transcription readthrough (Fold change) for the indicated Wdr82-HA tagged clonal cell lines was analyzed by RT-qPCR as in Figure 7F. Control, Wdr82-HA tagged cell line untransfected with the PNUTS expression plasmid. D, transcription readthrough defects in WT versus Wdr82-HA tagged (Control) cell lines.

**Figure S14**

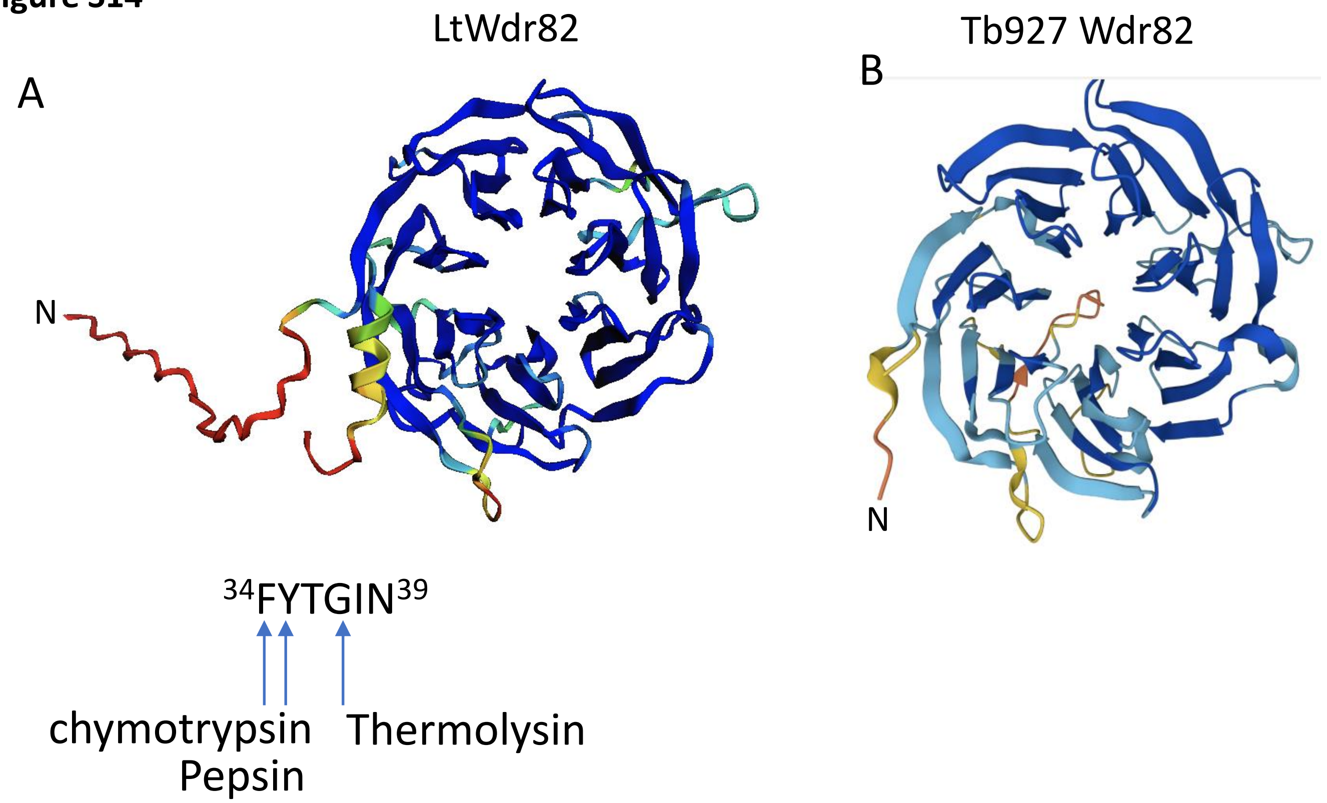


**Figure S14.** AlphaFold model of LtWdr82 has an unstructured N-terminus. AlphaFold model of (A) LtWdr82, and (B) TbWdr82. The structures are colored by pLDDT values, with blue representing high model confidence (pLDDT > 90) and orange representing low model confidence (70 > pLDDT > 50). Potential protease cleavage sites in the N-terminus of LtWdr82 are indicated. N, N-terminus.
